# Mutation rate and spectrum in *Caenorhabditis elegans* mutation accumulation lines subjected to RNAi-induced knockdown of the mismatch repair gene *msh-2*

**DOI:** 10.1101/2021.07.12.452034

**Authors:** Vaishali Katju, Anke Konrad, Thaddeus C. Deiss, Ulfar Bergthorsson

## Abstract

DNA mismatch repair (MMR), an evolutionarily conserved repair pathway shared by prokaryotic and eukaryotic species alike, influences molecular evolution by detecting and correcting mismatches that escape DNA polymerase proofreading, thereby protecting genetic fidelity, reducing the mutational load, and preventing lethality. Herein we conduct the first genome-wide evaluation of the alterations to the mutation rate and spectrum under impaired activity of the *MutSα* homolog, *msh-2*, in *Caenorhabditis elegans*. We performed mutation accumulation (MA) under RNAi-induced knockdown of *msh-2* for 50 generations in obligately outcrossing *fog-2(lf)* lines, followed by next-generation sequencing of 19 MA lines and the ancestral control. *msh-2* impairment substantially increased the frequency of nuclear base substitutions (∼23×) and small indels (∼328×) relative to wildtype. However, we observed no increase in the mutation rates of mtDNA, and copy-number changes of single-copy genes. There was a marked increase in copy-number variation of rDNA genes under MMR impairment. In *C. elegans*, *msh-2* repairs transitions more efficiently than transversions as well as increases the AT mutational bias relative to wildtype. The local sequence context, including sequence complexity, G+C-content, and flanking bases influenced the mutation rate. The X chromosome had a lower substitution and higher indel rate than autosomes, which can either result from sex-specific mutation rates or a nonrandom distribution of mutable sites in the genome. Comparison of MMR impairment in *C. elegans* to that in other species shows that the specificity of the MMR varies between taxa, and is more efficient in detecting and repairing small indels in eukaryotes relative to prokaryotes.

## INTRODUCTION

Mutation is a ubiquitous process across organisms, introducing novel genetic variation from which adaptive processes can sample. However, when mutational events facilitated by replication errors, exogenous (UV radiation, chemicals) or endogenous (oxygen free radicals) influences, are left entirely untempered, genome integrity falls victim to this same evolutionarily essential process (Lindahl et al. 1993; Kunkel and Erie 2005). A number of DNA repair mechanisms have evolved that limit the degenerative processes of DNA damage and replication errors. One such system is DNA mismatch repair (MMR), which corrects errors arising via base-base mismatches and small loop-outs resulting in indel mutations (Harfe and Jinks-Robertson 2000; Aquilina and Bignami 2001; Kunkel and Erie 2005; Jiricny 2006) by detecting the mismatch, differentiating between the parent and newly synthesized strand, and initializing repair (Modrich 1991; Kolodner 1996). MMR influences molecular evolution in various ways, most notably by detecting and correcting mismatches that escape DNA polymerase proofreading, thereby protecting genetic fidelity, reducing the mutational load and preventing lethality. Conversely, loss-of-function mutations in MMR genes can increase the mutation rate by several orders of magnitude. Furthermore, the specificity of the MMR system, with respect to the efficiency of MMR in recognizing and repairing different kinds of mismatch errors, contribute to the evolution of the base composition and structure of genomes.

MMR systems are found in functionally conserved form across the tree of life, consistent with their essentiality and ancient origin (Kolodner 1996; Eisen and Hanawalt 1999; Kunkel and Erie 2005; Lin et al. 2007; Sachadyn 2010). The *E. coli* (and other alpha-, beta-, and gamma-proteobacteria) MMR pathway contains a single pair of MutS and MutL proteins, while many other bacterial and eukaryotic genomes code for a repertoire of several MutS and MutL homolog proteins (MSH and MLH, respectively). Eukaryotic genomes encode several different MSH proteins, the sum of which perform a highly conserved MMR recognition mechanism (Groothuizen and Sixma 2016). MutS is required for the recognition of DNA lesions and recruits MutL for subsequent repair, which, in heterodimeric forms of their respective homologs, carry out the various functions of MMR (Modrich 1991; Harfe and Jinks-Robertson 2000; Kunkel and Erie 2005; Lin et al. 2007; Sachadyn 2010). However, the exact number of homologs and their exact functional complementation is not perfectly conserved (Denver et al. 2005; Groothuizen and Sixma 2016). *msh-4* and *msh-5* play important roles in chiasma formation during meiotic recombination (Shodhan et al. 2014), but have no known functions related to MMR (Zalevsky et al. 1999). The MSH2/MSH3 heterodimer (MutS*β*) corrects small and large loop-outs (no base-base mismatch repair) in yeast, humans and other mammals (Alani 1996; Habraken et al. 1996; Aquilina and Bignami 2001; Harrington and Kolodner 2007; Lee et al. 2007); yet *msh-3* has not been detected in several other metazoans, including *Drosophila* and *Caenorhabditis* (Tjisterman et al. 2002; Lin et al. 2007). Certain plants (*Arabidopsis thaliana*) form a MSH2/MSH7 (MutS*γ*) complex which corrects certain mismatches and loop-outs (Culligan and Hays 2000; Gomez and Spampinato 2013); however, the *msh-7* homolog has so far only been detected in plants. In contrast, the MSH2/MSH6 (MutS*α*) heterodimer complex is more strongly conserved across metazoan taxa (Lin et al. 2007). The MutS*α* complex is responsible for base-base mismatch and small loop-out indel detection, as well as the initiation of the repair cascade (Harfe and Jinks-Robertson 2000; Aquilina and Bignami 2001; Kunkel and Erie 2005; Richman 2015). A defect in, or insufficient concentration of the MutS*α* complexes results in microsatellite and genome instability and promotes tumorigenesis in humans (Aquilina and Bignami 2001; Degtyareva et al. 2002; Tijsterman et al. 2002; Lynch et al. 2009; Richman 2015; Meier et al. 2018).

Defects in the MutS*α* complex have been shown to lead to increased repeat tract length instability (mutation rate fold changes between 100 and 700) in multiple species, such as *S. cerevisiae* and *C. elegans* (Strand et al. 1993, Degtyareva et al. 2002; Tijsterman et al. 2002; Denver et al. 2005). Additionally, base substitution rates increased between five and more than 30-fold between wildtype and knockout lines of the MutS*α* complex (Yang et al. 1999; Degtyareva et al. 2002; Tijsterman et al. 2002; Denver et al. 2005). However, the majority of these studies did not consider the genome as a whole, but rather relied on mutation rate estimates based on partial genome analysis or on specific reporter genes. For example, the Denver et al. (2005) analysis was restricted to 20 kb, which represents less than 0.02% of 100.3 Mb *C. elegans* genome. Differences in mutational rates and spectra arising due to sequence context, genome architecture, functional content, and transcription or replication cannot be captured in approaches that do not evaluate the genome as a whole. Mutation accumulation (MA) experiments coupled with whole genome sequencing (WGS) provides an experimental framework to assess such variability (reviewed in Katju and Bergthorsson 2019). MA experiments consist of passaging independently evolving populations through single individual bottlenecks, thus rendering the effects of natural selection minute. All but the most deleterious or lethal mutations are thus allowed to accumulate as if neutral with respect to fitness, thereby facilitating a more comprehensive assessment of the mutational process. The first such experiments utilizing *msh-2* knockout mutant lines followed by next-generation sequencing in the unicellular yeast *S. cerevisiae* (Lang et al. 2013) and the angiosperm model plant *A. thaliana* (Belfield et al. 2018) demonstrated an estimated 225-fold and >1000-fold increase in the spontaneous mutation rate, respectively.

Although several preceding studies in *C. elegans* have offered insights into the mutational consequences of *msh-2* deficiency, they have been limited in their scope owing to a narrow focus on specific reporter genes and known mutational hotspots, or on a limited number of nuclear and mitochondrial loci across the genome (Degtyareva et al. 2002; Tijsterman et al. 2002; Denver et al. 2005). Mutation accumulation (MA henceforth) studies with *C. elegans msh-2* knockout mutants are challenging as they have greatly reduced fertility and *msh-2* lines that are maintained by single individual descent typically become extinct in 10-20 generations (Degtyareva et al. 2002). In lieu of *msh-2* knockout mutation lines, we employed a *msh-2* RNA interference (RNAi henceforth) approach to knockdown the expression of *msh-2* during the MA process. Although a knockdown may not capture the full effect of a gene knockout, it enables a more extended MA experiment which helps illuminate the kinds of mutations that accumulate with an MMR system defective in *msh-2.* Several studies have shown differences in mutation rates between males and females which can manifest as different mutation rates between the X chromosome and the autosomes. In addition to the RNAi knockdown approach, we used an obligately outcrossing (male-female) line of *C. elegans* owing to a mutation in the *fog-2* gene. Although these deviate from the normal reproductive mode of *C. elegans*, which is predominantly self-fertilizing hermaphroditism, the results facilitate a comparison of the number of mutations between the X chromosome and the autosomes as males are XO and females are XX. A higher mutation rate in males therefore predicts a higher mutation rate on the autosomes than on the X chromosome (Miyata et al. 1987). These experiments provide the first genome-wide view of the MA process in *C. elegans* with an impaired *msh-2* and present an analysis of the greatest number of mutations in any MMR-deficient background in the species.

## MATERIALS AND METHODS

### Mutation accumulation (MA) lines subjected to msh-2 knockdown

The MA lines used in the *msh-2* knockdown experiment were established from a single male-female pair of the obligatory outcrossing *fog-2(lf)q71* mutant strain of *C. elegans* (Schedl and Kimble 1988; Katju et al. 2008; Farslow et al. 2015). Following four generations of single-pair sib-mating, two males and one female from the F_5_ offspring of the founding pair were used to establish 74 MA lines (**Supplemental Figure S1**) (Katju et al. 2008). The remaining founder population was frozen at -80° C to be used as the ancestral control, referred to as the pre-MA ancestral control (PMAC). The MA lines were subjected to repeated population bottlenecks by randomly picking one female and two male L4 larval worms to establish the subsequent generation (**Supplemental Figure S1**). Given a population bottleneck of three individuals (two males and one female) each generation, the effective population size, *N_e_*, is approximately 2.67 individuals (Wright 1931). Hence, the *N_e_* of this MA experiment was similar to that of our MA experiment with obligately selfing lines maintained at population bottlenecks of *N* = 1 per generation (*N_e_* = 1; Katju et al. 2015, 2018). Under both these MA regimes, mutations with selection coefficients less than approximately 10% are expected to contribute to mutational degradation given that they will accumulate within these MA lines at the neutral rate, although they may not necessarily neutral with respect to absolute fitness. However, our previous analyses of the accumulation of SNPs and small indels in *C. elegans* MA lines found no differences between MA lines due to *N_e_* (Konrad et al. 2019).

The expression of the *msh-2* mismatch repair (MMR) gene was knocked down each MA generation via a standard RNAi feeding protocol (Kamath et al. 2001) in order to elevate the accumulation of germline and somatic mutations (Degtyareva et al. 2002; Tijsterman et al. 2002). The gene of interest, *msh-2*, was amplified via PCR and cloned into the L4440 vector, which has two T7 promoters in inverted orientation flanking the cloning site. The cloned plasmids were transformed into HT115(DE3), an RNase III-deficient *E. coli* strain with IPTG-inducible expression of the T7 polymerase. A bacterial clone containing the feeding vector with the *msh-2* gene was obtained from Julie Ahringer at the University of Cambridge. Single colonies of HT115 bacteria containing cloned L440 plasmids with the *msh-2* gene were picked and cultured in LB with 50 μg/ml ampicillin for approximately 8-12 hours. In order to induce the expression of dsRNA of the *msh-2* gene, these cultures were then seeded directly onto NGM plates with 1mM IPTG and 50 μg/ml ampicillin. Seeded plates were allowed to dry at room temperature and the induction of dsRNA was continued overnight. For each generation of MA, one female and two male siblings were placed onto the freshly prepared NGM feeding plates containing seeded bacteria expressing dsRNA for the *msh-2* gene. The resulting progeny were maintained on these RNAi feeding plates for four days at 20°C, following which another generation of MA on new RNAi feeding plates was initiated in the manner described above. To prevent accidental losses of the experimental lines, plates from the preceding four generations were maintained as backups in a separate 20°C chamber. Each line was subjected to 50 generations of MA, with bottlenecking and RNAi treatment at each generation. MA lines that went extinct over the course of the experiment were re-initiated from populations of the previous generations up to five times prior to being considered formally extinct. Four lines became extinct during this mutation accumulation phase due to a complete lack of recruitment of new generations of individuals, despite backup attempts from preceding generations.

It is critical that mutations accumulated in the MA phase of the experiment are fixed within each line and not capable of segregation as wildtype alleles. To achieve this, each extant MA line was subjected to fifteen additional generations of full-sibling mating without RNAi treatment. Treating the last MA generation as the reference population. In practice, the reduction in heterozygosity should be greater than given that the MA treatment involved a strict regime of full-sib mating, even though a single female was paired with and could be multiply-mated to two brothers. Thereafter, all surviving MA lines (total 70 lines) were frozen at -80°C. Nineteen of the extant 70 MA lines and the ancestral control were sequenced for this study. This set included five MA lines previously observed to have experienced the most extreme decline in fitness (Farslow et al. 2015), and 14 additional randomly chosen lines.

### Genomic DNA extraction, whole-genome sequencing, and alignment

A total of 20 *C. elegans fog-2* lines comprising the ancestral control and 19 *msh-2* knockdown MA lines were prepared for DNA whole-genome sequencing. For each line, a trio of one female and two male worms were used to establish a population that was expanded for two to three additional generations to generate sufficient worm tissue for genomic DNA (gDNA) extraction. Sequencing followed the methodologies previously described (Konrad et al. 2017, 2018, 2019). The PureGene Genomic DNA Tissue Kit (QIAGEN no. 158622) and a supplemental nematode protocol were used for isolation of gDNA. DNA quality and concentration were checked on 1% agarose gels through electrophoresis, a Nanodrop spectrophotometer (Thermo Fisher), and BR Qubit assay (Invitrogen). Sonication of 2*μg* of each DNA sample in 85*μL* TE buffer yielded target fragment lengths of 200-400 bp, which were end-repaired (NEBNext end repair module (New England BioLabs)) and purified (Agencourt AMPure XP beads (Beckman Coulter Genomics)). Beads were removed after adapter ligation as previously described (Thompson et al. 2013). Custom pre-annealed Illumina adapters were utilized and ligated to the purified fragments. 3’ adenine overhangs were added (AmpliTaq DNA Polymerase Kit, Life Technologies). PCR amplification was performed via Kapa Hifi DNA Polymerase (Kapa Biosystems) and Illumina’s paired end genomic DNA primers containing 8 bp barcodes. Size fractionation of the PCR products was performed on 6% PAGE gels and 300-400bp fragments were selected. Gel extraction by diffusion at 65°C and gel filtration (NanoSep, Pall Life Sciences) followed by Agencourt AMPure bead purification was used to generate the final fully purified fragments. The fragments’ final quality and quantity was checked via the Agilent HS Bioanalyzer and HS Qubit assays. Multiplexed libraries were sequenced on Illumina HiSeq sequencers with default quality filters at the Northwest Genomics Center (University of Washington).

The demultiplexed raw reads were aligned to the reference N2 genome (version WS247; www.wormbase.org; Harris et al. 2010) using the Burrows-Wheeler Aligner (BWA Version 0.5.9) (Li and Durbin 2009) and Phaster (Green lab) (previously described in Konrad et al. 2018, 2019).

### Sequence alignment and identification of putative single nucleotide polymorphisms and indels

The identification of small mutations (SNPs and indels up to 100 bp long) followed the same methodology previously used to identify mutations in the wildtype MA lines relative to their ancestral control (Konrad et al. 2019). Briefly, alignment files from Phaster and BWA were analyzed separately. Putative SNPs and indels were identified using Platypus (Rimmer et al. 2014), Freebayes (Garrison and Marth 2012), and a pipeline consisting of mpileup (Li et al. 2009), bcftools (Li 2011), vcfutils (Danecek et al. 2011) and custom filters written in Perl. All putative variants were filtered against the ancestral control line, and Indelminer (Ratan et al. 2015) was used as an additional approach to call indels with the ancestral control genome as a direct reference (normal sample). Putative indel identification was primarily based on the Phaster alignments due to its greater ability to split reads and align with gaps. However, BWA alignments were used to verify these indel calls. A minimum root-mean-square mapping quality of 30 and 40 was required for SNPs and indels to be retained, respectively. A minimum of three and five high quality reads were required to support each SNP and indel, respectively. Putative variants present in the ancestral control genome, even with low-quality or coverage, were removed from the analysis. A minimum of 80% of all high-quality calls at the variant position were required to support a variant in question for it to be retained in the dataset. Finally, each variant had to be called independently by at least two of the variant callers in order to be considered for further analysis.

To rule out putative variants identified due to sequencing or alignment error, every variant was independently verified by calculating a binomial probability for it, given the number of variant calls at the same location in the genome across all other populations sequenced (Konrad et al. 2019). For each putative variant position, the number of reads across all lines calling the variant were summed and divided by the total number of reads at the variant position. We used this as the probability of any given read calling the variant by chance (*P*). For each putative mutation, we counted the number of reads within every individual line which called the variants (*K*), and the total number of reads at the position in that line (*N*). We then calculated the *p*-value for the variant (*var*) in that line 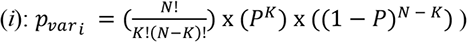. The *p*-values across all lines were sorted from most significant to least significant, and a Holm-Bonferroni correction was applied to determine if the variants called by the previous pipeline met the critical *p*-value threshold.

### Annotation, characterization, and mutation rate calculation for SNPs and small indels

All identified variants were annotated using a custom script and the GFF file available for the N2 reference genome of *C. elegans* (version WS247; www.wormbase.org; Harris et al. 2010). Mutations were assigned to exons, introns, and intergenic regions, as well as to chromosomal arms, cores, and tips based on boundaries predicted by Rockman and Kruglyak (2009). The tips domains contain roughly 7% of the *C. elegans* genome and have high gene density and extremely low recombination rates. The arms domains (26%) have high recombination rate and the center domains (cores, 47%) have low recombination rates. The boundaries in the original analysis of recombinational domains were identified using segmented linear regression (Rockman and Kruglyak 2009). Mutation rates (μ_*var_i_*_) were estimated individually for each line and across genomic subdivisions as variants per base per generation 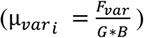, where *F_var_* equals the number of substitutions or indels within the line, *G* refers _-∗/_ to the number of generations, and *B_total_* refers to the number of bases in the genome or genomic subdivision that meet the same thresholds as for variant identification (version WS247). For mitochondrial mutation rates, the frequencies of variants were calculated as a percentage of quality reads calling the variant. Overall mutation rates were calculated by averaging the line-specific mutation rates within: 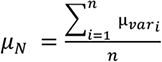, where *var_i_* refers to the line-specific mutation rate, and *n* refers to the total number of lines. The number of MA generations through which each population was propagated differed between the lines (**Supplemental Table S1**).

Genomic repeat regions and homopolymeric runs were identified using the Imperfect Microsatellite Extractor (IMEX 2.1; Mudunuri and Nagarajaram 2007). Homopolymeric runs of at least 6 bp in length were included. For di-, and tri-nucleotide repeats, we required at least four repetitions of the motif, while three repeats of each individual repeat unit were required for tetra-, penta-, and hexa-nucleotide repeats. Imperfect repeats were not included in the analysis of repeats, unless the imperfect repeat divided the overall repeat region into at least one run that met the above criteria. Every putative variant was mapped against this final list of genome-wide repeats.

Every protein-coding gene was categorized as either a germline or non-germline expressed gene based on the data of Wang et al. (2009). The mutation rate in germline expressed genes was calculated by summing the number of mutations within each line that mapped to germline expressed genes and dividing by the total number of high-quality bases within germline expressed genes. Mutation rates for non-germline expressed genes were calculated in the same fashion.

Sequence complexity was calculated as described in Morgulis et al. (2006). Briefly, given a sequence (*a*) of length *n* and 64 possible triplets of {A, C, G, T}, the occurrence of each possible triplet (*t*) was counted across the sequence and yields *c_t_(a)*. The total number of overlapping triplets occurring in any sequence (*l*) equals *n*-2. Sequence complexity (*S(a)*) was then calculated as:

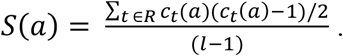

All statistical tests were performed in R (R Core Development Team 2014).

### Genomic properties influencing mutation rates

A regularized logistic regression approach was used to determine the genomic properties most indicative of the mutability of different sites in the genome (Ness et al. 2015; Konrad et al. 2019). The training set for the modeling consisted of 1,000,000 random non-mutated sites throughout the *C. elegans* genome combined with the 3,125 unique substitution sites. Chromosomal location, functional properties, germline expression (Wang et al. 2009), recombination rate (Rockman and Kruglyak 2009), G+C-content and sequence complexity (*s*; Morgulis et al. 2006), repeat sequence, chromatin state (Evans et al. 2016), periodic A_n_/T_n_-clusters (PATCs; Frøkjær-Jensen et al. 2016), and trinucleotide sequence context for each of the 1,003,125 sites (approximately 1% of the genome, each) were compiled as potential predictors for mutability (Konrad et al. 2019). G+C-content and sequence complexity (*s*) were calculated for 41 bp windows around each site (Konrad et al. 2019). All categorical predictors (chromosome, functional category, trinucleotide context) were converted to a series of binary predictors referring to each category level. Recombination rate, G+C-content, and sequence complexity were treated as numeric predictors, while germline expression and repeat sequence were binary predictors.

The GLMnet package (v1.9-8) was used to perform a generalized linear model fit in R (R Core Development Team 2014). This package implements penalized maximum likelihood through ridge and lasso regression, resulting in more precise fits for models which are built with intercorrelated predictor variables (Friedman et al. 2010). The response variable was binary: 1 for a mutation and 0 for a random site/no mutation. The penalty against significant correlations between predictor coefficients was determined by the regularization parameter (λ). λ was set to 6.83 × 10^-5^, which is the value (lambda.min) at which the cross-validated error is minimized through the built-in cross-validation function during the model building step. An *α* of 0.01 was used to retrieve the model coefficients, which shrinks correlated predictor coefficients together. Varying *α* did not affect the model fit. Odds ratios (OR) for predictors were calculated as OR = *e^c^*, where *c* refers to a given predictor coefficient.

For each site in the genome, its mutability was estimated as the probability of encountering a mutation, using the predict function of GLMnet with the model coefficients estimated above for each genomic predictor. The probabilities of mutation at any given site are affected by the proportion of mutated sites over random sites used during the model training step. Hence, we are interested in the relative mutability values. Given the 3,125 SNP sites (or 9,858 unique indel sites) and 1,000,000 non-mutated sites, the mean predicted mutability is approximately 0.002. The predicted mutability across the genome ranged from 0.75 × 10^-4^ to 0.57. 100% of SNPs were covered between mutabilities of 0.0 and 0.68 for SNPs. Mutabilities from 0 to 0.12 (encompassing > 99.9 % of SNPs and >99.9 % of genomic sites) were combined into bins of size 0.015, and mutation rates were calculated for each 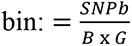, refers to the number of SNPs per bin, *B* refers to the number of sites in each bin, and *G* refers to the average number of generations (40.3). Correlation coefficients and *R*^2^ values were calculated for a linear (Pearson) regression of mutation rate over mutability in R (R Core Development Team 2014).

### Data Availability Statement

Whole-genome sequence data of all experimental lines and the ancestral control have been deposited at the National Center for Biotechnology Information Sequence Read Archive (Bioproject PRJNA 554105).

## RESULTS

We sequenced the genomes of the ancestral control and 19 *fog-2* MA lines subjected to *msh-2* knockdown with an average read depth of 30.35× and 16.51×, respectively (**Supplemental Figure S1**; **Supplemental Table S1**). The proportion of the genome included for SNP analysis across the experimental *msh-2* knockdown MA lines ranged from 95−97%. More stringent filtering thresholds imposed for the indel analysis led to the inclusion of 59−69% of the genome. The whole-genome sequences of the MA lines can be accessed through the National Center for Biotechnology Information Sequence Read Archive (Bioproject PRJNA554105).

Prior to combining the mutation rate data for the five lowest fitness MA lines with the additional randomly chosen 14 MA lines, we tested if there was a difference in the base substitution and the indel rates between the two groups. The average base substitution rate for the five low fitness and 14 randomly chosen MA lines was 3.99 × 10^-8^ /site/generation and 4.30 × 10^-8^ /site/generation, respectively. The slightly higher value for the randomly chosen MA lines was not significantly different from the lines with the lowest fitness (*t* = 0.89, *p* = 0.38). The results for small indels were very similar. The small indel rate for the low fitness and the randomly chosen MA lines was 2.02 × 10^-7^ /site/generation and 2.31 × 10^-7^ /site/generation, respectively. The difference in indel rates was not significant (*t* = 1.54, *p* = 0.14). Given that the two mutation rates are essentially equivalent, we used the combined dataset of 19 MA lines for all downstream analyses.

### Significantly elevated nuclear mutation rates in msh-2 knockdown MA lines relative to spontaneous MA lines

The vast majority of mutations identified in the *msh-2* MA lines were small indels and single nucleotide substitutions in the nuclear genome. We identified 3,125 substitutions and 10,861 small indels across the 19 *msh-2* knockdown MA lines following an average of only 40.3 MA generations, resulting in a combined mutation rate of 2.65 × 10^-7^ mutations per site per generation. This yields a nuclear substitution and indel rate of 4.22 (95% CI: ± 0.30) × 10^-8^ and 2.23 (95% CI: ± 0.17) × 10^-7^ mutations per site per generation, respectively (**Table 1**; **Figure 1A**). We compared these mutation rates to our preceding analyses of spontaneous MA lines of *C. elegans* maintained at population bottlenecks of *N* = 1 individual per generation (Konrad et al. 2019). We henceforth refer to the spontaneous MA lines as wildtype as they represent the spontaneous mutational input in the absence of selection but under a functional DNA-repair regime. Relative to wildtype, the mutation rates of the *msh-2* MA lines exhibit a ∼23-fold and a ∼328-fold increase in the base substitution and small indel rate, respectively (**Table 1**; **Figure 1B**). Relative increase in mutation rate in *msh-2* knockdown MA lines compared to wildtype MA lines is significantly greater for small indels compared to SNPs (*t*-test: *t =* 24.43, *p* = 1.61 × 10^-15^). The significantly higher frequency of indels relative to substitutions in the *msh-2* knockdown lines (*t*-test: *t =* -20.92, *p =* 1.18 × 10^-14^) translates into a ratio of 3.85 indels per substitution (**Figure 1C**), which is a significantly higher ratio than that observed in the wildtype lines (0.32 indels per SNP; **Figure 1C**; *t*-test: *t =* 12.92, *p =* 6.92 × 10^-10^).

**Figure 1.**
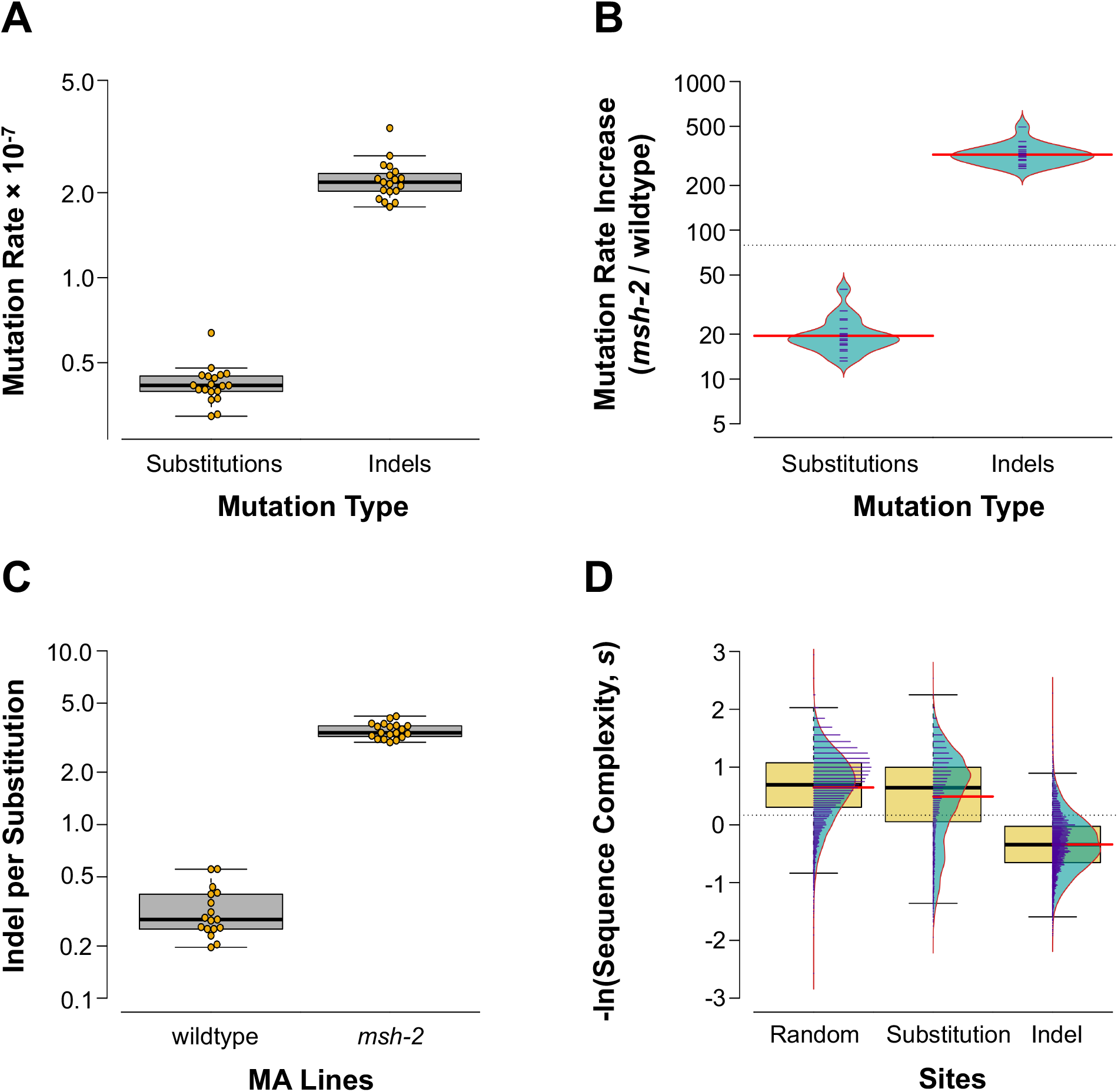
Increase in mutation rates in *msh-2* knockdown MA lines compared to wild-type. *(A)* Base substitution rates are significantly lower than small indel rates in the *msh-2* knockdown lines. Rates are calculated as per site per generation. *(B)* Mutation rate increase in *msh-2* knockdown MA lines compared to wildtype MA lines is significantly greater for small indels compared to SNPs. *(C)* The indels/substitution ratio is significantly higher in the *msh-2* knockdown MA lines compared to wildtype MA lines. *(D)* Sequence complexity in the vicinity of indel mutations is significantly lower than that for SNPs or randomly chosen genomic sites. Sequence context in the vicinity of SNPs is significantly lower than that for random sites. Thick horizontal black lines and red lines indicate the sample median and mean, respectively.

**Table 1.** Mutation rates (*µ* × 10^-7^) and fold-changes in *msh-2* MA lines (this study) in comparison to spontaneous MA lines of the laboratory *N2* Bristol strain (wildtype) of *C. elegans* (Konrad et al. 2017, 2018, 2019). Substitution (SNP), small insertion and deletion (Indel), small deletion (Del) and insertion (Ins), and mitochondrial (mt) rates are given as mutations per site per generation. Copy-number gains (CNV+) and losses (CNV-) are listed as changes per protein-coding gene per generation. Line 1T of the wildtype lines was not included as it included unusually large copy-number changes indicative of extensive chromosomal rearrangements (Konrad et al. 2018). Statistically significant differences in rates between the two data sets are indicated by asterisks. *wt* refers to wildtype.

| Mutational Class | *wt* ($\mu \times 10^{-7}$) | *msh-2* ($\mu \times 10^{-7}$) | $\mu_{msh-2} / \mu_{wt}$ | Significance |
| --- | --- | --- | --- | --- |
| $\mu_{\text{SNP}}$ | 0.0184 | 0.42 | 22.93 | * |
| $\mu_{\text{Indel}}$ | 0.0068 | 2.23 | 327.94 | * |
| $\mu_{\text{Del}}$ | 0.0051 | 1.47 | 288.24 | * |
| $\mu_{\text{Ins}}$ | 0.0018 | 0.76 | 422.22 | * |
| $\mu_{\text{mt}}$ | 1.05 | 2.18 | 2.08 | — |
| $\mu_{\text{CNV+}}$ | 26.40 | 10.70 | 0.41 | — |
| $\mu_{\text{CNV-}}$ | 11.90 | 3.95 | 0.33 | — |

Low complexity DNA sequence repeats have a significant effect on the mutation pattern (Morgulis et al. 2006; **Figure 1D**; ANOVA: *F =* 9117, *p* < 2 × 10^-16^). The complexity in the vicinity of the small indels was significantly lower than that associated with either base substitutions (**Figure 1D**; Tukey’s Multiple Comparison: *p =* 0.00; *t*-test: *t =* 59.12, *p* < 2.2 × 10^-16^) or with random sites in the genome (Tukey’s Multiple Comparison: *p =* 0.00; *t*-test: *t =* 136.99, *p* < 2.2 × 10^-16^). Moreover, the sequence context around base substitutions was less complex than that observed in the vicinity of random sites (Tukey’s Multiple Comparison: *p =* 0.00; *t*-test: *t =* 10.75, *p* < 2.2 × 10^-16^).

### Base substitutions accumulate neutrally within exons

The synonymous [3.97 (95% CI: ± 0.70) × 10^-8^ /site/generation] and nonsynonymous [4.46 (95% CI: ± 0.36) × 10^-8^/site/generation] substitution rates were not significantly different from one another in the *msh-2* MA lines (**Supplemental Figure S2A**; *t*-test: *t* = 1.21, *p* = 0.24). Furthermore, the relative increase in synonymous and nonsynonymous substitutions in the *msh-2* MA lines relative to wildtype MA lines were not significantly different (*t*-test: *t* = 1.59, *p* = 0.13). Although the average nonsynonymous/synonymous substitution ratio (*K_a_/K_s_*) for the *msh-2* MA lines appeared to be lower than that of N2 wildtype MA lines, there was no significant difference between them (*t*-test: *t* = 1.23, *p* = 0.23). Furthermore, the average *K_a_/K_s_* in the *msh-2* MA lines was not significantly different from unity, which is consistent with negligible purifying selection in exons during the MA experiment (*t*-test: *t* = 1.83, *p* = 0.09). Frameshift mutations [1.04 (95% CI: ± 0.09) × 10^-8^/site/generation] were less frequent than either synonymous (*t*-test: *t* = 6.00, *p* = 1.00 × 10^-5^) or nonsynonymous substitutions (*t*-test: *t* = 18.03, *p* = 5.09 × 10^-14^) (**Supplemental Figure S2A**). However, the increase in the rate of frameshift mutations (50×) in the *msh-2* MA lines compared to wildtype was significantly greater than the increase in either the synonymous (29×; *t*-test: *t* = 5.98, *p* = 7.75 × 10^-7^) or nonsynonymous (24×; *t*-test: *t* = 10.08, *p* = 3.08 × 10^-10^) substitution rate (**Supplemental Figure S2B**).

### Substitution bias in msh-2 knockdown MA lines

Transitions outnumbered transversions in *msh-2* knockdown MA lines, while the opposite was observed in the wildtype MA lines (Konrad et al. 2019). Consequently, the transition to transversion ratio was significantly higher for *msh-2* knockdown (mean = 1.15) than for wildtype (mean = 0.67) MA lines (**Figure 2A**; *t*-test: *t =* 7.69; *p =* 6.84 × 10^-9^). There was a lower mutational bias towards A+T during *msh-2* knockdown than in the wildtype MA lines (**Figure 2A**; *t*-test: *t =* 7.36, *p =* 8.75 × 10^-8^). In the *msh-2* knockdown MA lines, 64% of the base substitutions that change the G+C-content are towards an increase in A+T-content, whereas in wildtype N2 MA lines, this proportion was 78%. The increase in substitution rates in the *msh-2* MA lines compared to wildtype varied significantly between substitution types (**Figure 2B**; ANOVA: *F* = 36.41, *p* < 2 × 10^-16^). The greatest increase in specific substitution rates between *msh-2* knockdown and wildtype MA lines were in A/T → G/C transitions (54×) and A/T → C/G transversions (46×). In contrast, G/C → C/G transversions increased the least (3×).

**Figure 2.**
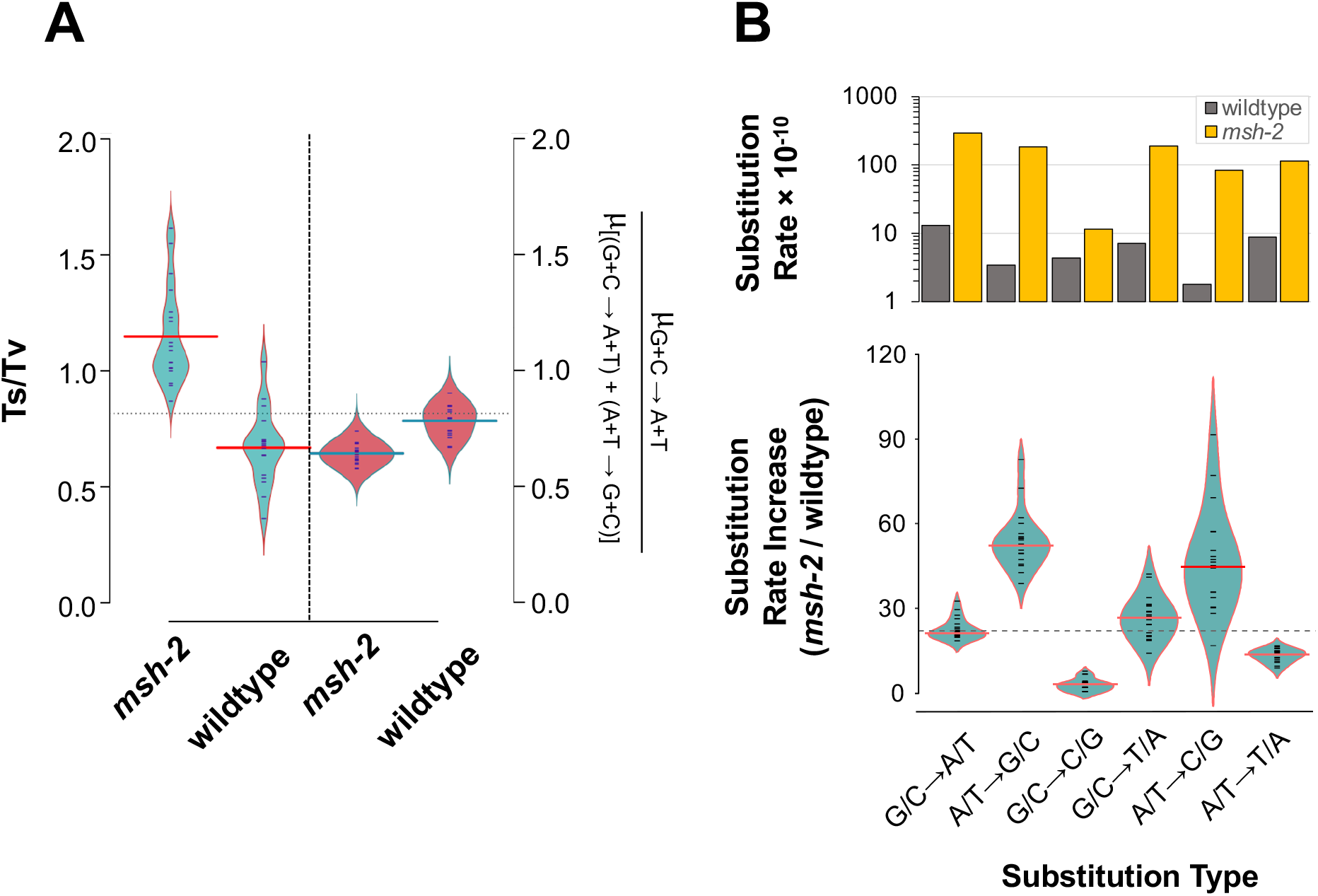
Changes in the mutational spectrum in *msh-2* knockdown versus wild-type MA lines. *(A)* Transition to transversion (Ts/Tv) ratios are significantly greater for *msh-2* knockdown than for wildtype line. *msh-2* knockdown lines exhibit a significantly lower G+C → A+T mutational bias relative to wildtype MA lines. *(B)* The frequency of A+T → G+C substitutions show a greater increase than G+C → A+T changes in the *msh-2* knockdown MA lines relative to wildtype.

### Sequence context influences mutation rates in the msh-2 knockdown MA lines

A/T → T/A transversions exhibited the greatest context-dependence (**Supplemental Figure S3**). While A/T → T/A transversions in both wildtype and *msh-2* knockdown MA lines occurred disproportionately between neighboring A and T nucleotides, the exact context-dependence differed between the two experiments. In the *msh-2* lines, the strongest context-dependence was for mutations between 5′−A<u>A</u>T−3′ and 5′−A<u>T</u>T−3′ (**Supplemental Figure S3A**; base of focal mutation is underlined), while it was strongest for mutations between 5′−T<u>A</u>A−3′ to 5′−T<u>T</u>A−3′ in the N2 wildtype lines (Konrad et al. 2019; **Supplemental Figure S3B**). There is a pronounced difference in the effects of *msh-2* knockdown on A/T → T/A transversions, between 5′−A and 3′−T on one hand (19-fold increase in exons, 152-fold increase in noncoding DNA) and 5′−T and 3′−A on the other (no increase in exons, 3-fold increase in noncoding DNA, **Supplemental Figure S3C**). The frequency of C/G → T/A transitions were especially increased in exons when the focal C was positioned between two cytosines (**Supplemental Figure S3C**). The same was true for A/T → C/G transversions when the focal A was positioned between a 5′−A and 3′−G context in both exons and noncoding DNA (**Supplemental Figure S3C**). In intergenic regions, this same substitution experienced a higher than usual increase in mutation rate when the focal A was flanked by a 5′−A and a 3′−non-A nucleotide (**Supplemental Figure S3C**).

A/T → T/A transversions were particularly common at the boundaries of homopolymeric runs of As and Ts. These types of homopolymeric runs are common in introns and intergenic regions, but not in exons. Furthermore, sequence complexity was significantly lower in the vicinity of A/T → T/A transversions relative to all other substitution types (**Table 2**; **Figure 3A**; ANOVA: *F* = 427.7, *p* < 2 × 10^-16^; Tukey’s Multiple Comparisons: *p* = 0.00 for all pairwise comparisons between A/T → T/A and other substitution types). 80% of all substitutions occurred in complex sequence. In contrast, only 18% of A/T → T/A transversions fell within complex sequence, while 82% of these mutations were adjacent to repetitive sequence (**Table 2**; **Figure 3B**). In fact, 72% of all substitutions falling within repeat sequences were A/T → T/A transversions (**Figure 2B**). These repetitive sequences are almost exclusively homopolymeric A or T runs that frequently flank A/T → T/A transversions. Finally, the frequency of SNPs adjacent to A and T homopolymeric runs was positively correlated with the length of the homopolymeric run (**Figure 3C**; Pearson Correlation Coefficient *r* = 0.93, *p* = 3.33 × 10^-5^).

**Figure 3.**
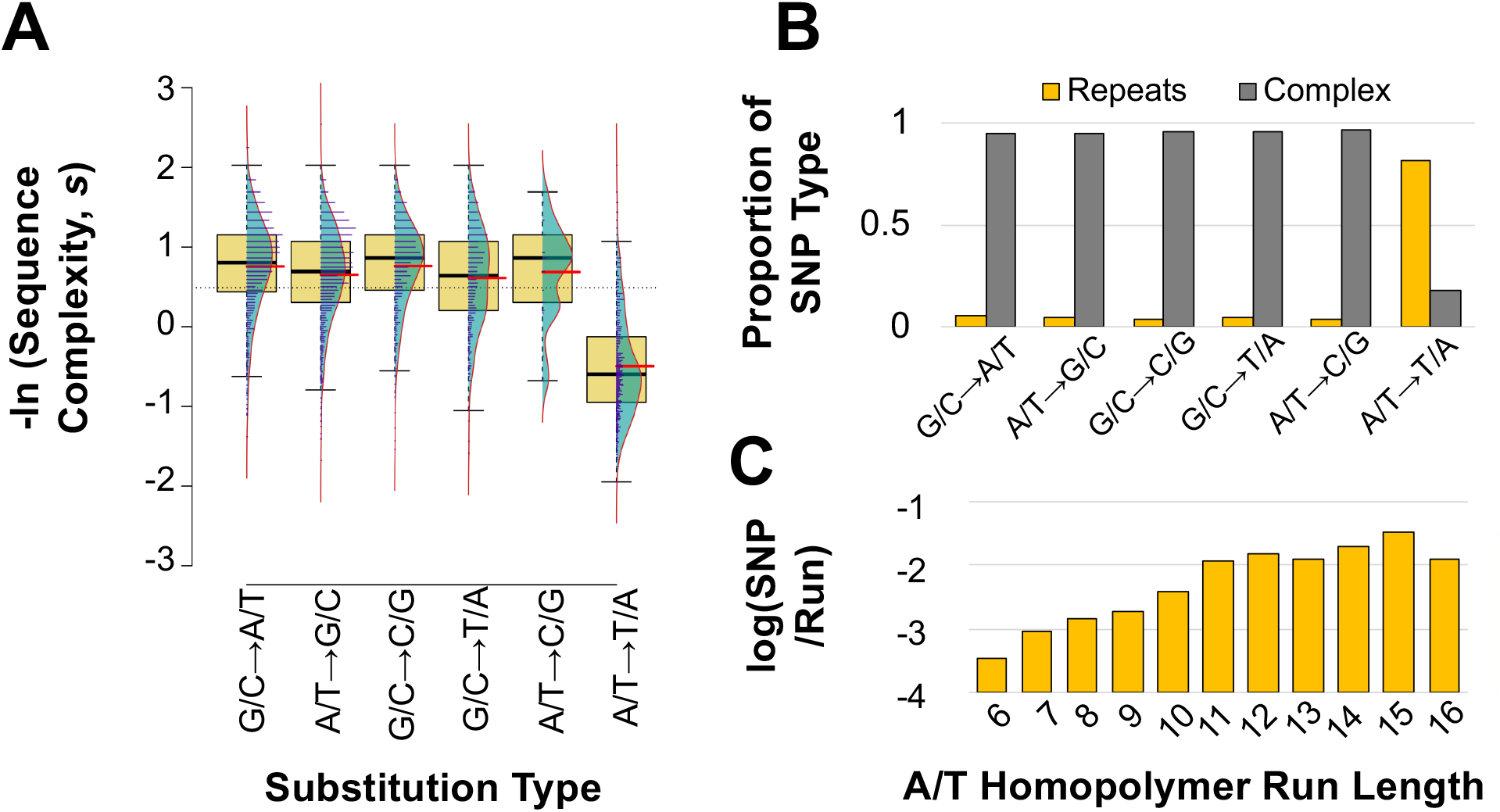
Relationship between sequence complexity and length of homopolymeric runs with individual substitution rates of *msh-2* knockdown MA lines. *(A)* Significantly lower DNA sequence complexity of regions flanking A/T→T/A substitutions relative to all other substitution types. Thicker horizontal black and red lines show the sample median and mean, respectively. *(B)* A/T → T/A substitutions are significantly more frequent in homopolymeric runs (length ≥ 6 bp) than other substitution types. *(C)* The number of A/T → T/A mutations per A or T repeat (normalized by number of respective repeats across the genome) are positively correlated with homopolymer length.

**Table 2.**
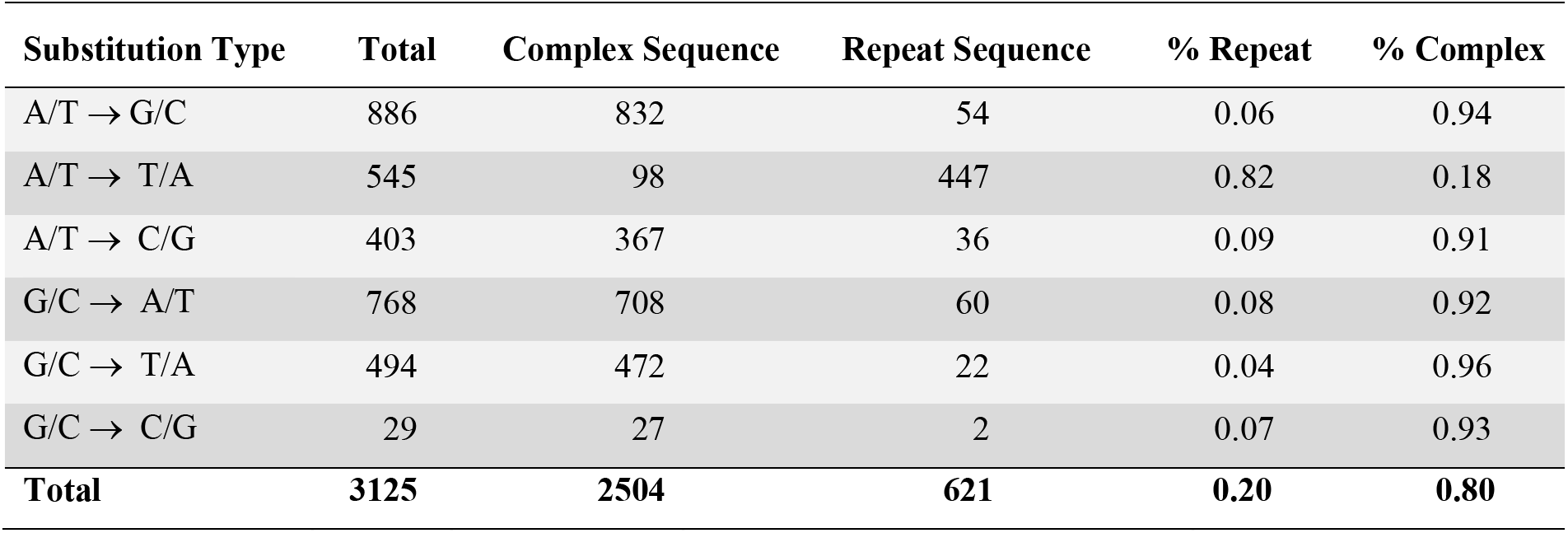
The majority of substitutions fall within complex sequence, with the exception of A/T → T/A substitutions, which are primarily within A+T-rich repeat regions of the genome.

| **Substitution Type** | **Total** | **Complex Sequence** | **Repeat Sequence** | **% Repeat** | **% Complex** |
| --- | --- | --- | --- | --- | --- |
| A/T → G/C | 886 | 832 | 54 | 0.06 | 0.94 |
| A/T → T/A | 545 | 98 | 447 | 0.82 | 0.18 |
| A/T → C/G | 403 | 367 | 36 | 0.09 | 0.91 |
| G/C → A/T | 768 | 708 | 60 | 0.08 | 0.92 |
| G/C → T/A | 494 | 472 | 22 | 0.04 | 0.96 |
| G/C → C/G | 29 | 27 | 2 | 0.07 | 0.93 |
| **Total** | **3125** | **2504** | **621** | **0.20** | **0.80** |

### Genomic heterogeneity in the substitution rate

The base substitution rate in exons, introns, and intergenic regions was 3.82 (95% CI: ± 0. 32), 4.60 (95% CI: ± 0.34), and 4.60 (95% CI: ± 0.42) × 10^-8^ substitutions/site/generation, respectively (**Figure 4A**). Despite the apparent low substitution rates in exons relative to noncoding DNA, the differences were not significant when their location in the genome was taken into account (3-way ANOVA: *F =* 2.19, *p =* 0.11). There is a significant interaction between the substitution rates in exons, introns, and intergenic regions with their location in cores and arms (**Figure 4B**; 3-way ANOVA: *F* = 2.75, *p* = 0.03). While there is no apparent difference based on coding content in chromosomal cores, the substitution rate in exons is 71% of that observed in noncoding DNA (introns and intergenic regions) in the chromosomal arms (**Figure 4B**). Coding content significantly influences the difference in substitution rates between *msh-2* and wildtype MA lines (**Figure 4C**; ANOVA: *F* = 6.86, *p* = 0.002). Exons and intergenic regions exhibit significantly greater increase in substitution rates relative to introns (exons versus introns: Tukey’s Multiple Comparisons, *p* = 0.002; intergenic versus introns: Tukey’s Multiple Comparisons, *p* = 0.027). However, there was no difference in the substitution rate increase observed in exons versus intergenic regions (Tukey’s Multiple Comparisons: *p* = 0.63). Higher substitution rates in noncoding DNA relative to exons is best attributed to the much higher frequency of context-dependent A/T to T/A transversions in the former (**Figure 4D**). These substitutions occur predominantly at the ends of homopolymeric runs of As and Ts, which are less common in exons. When A/T to T/A transversions are excluded from the analysis, there is no difference in the substitution rates between exons, introns, and intergenic regions (ANOVA: *F* = 0.08, *p* = 0.92).

**Figure 4.**
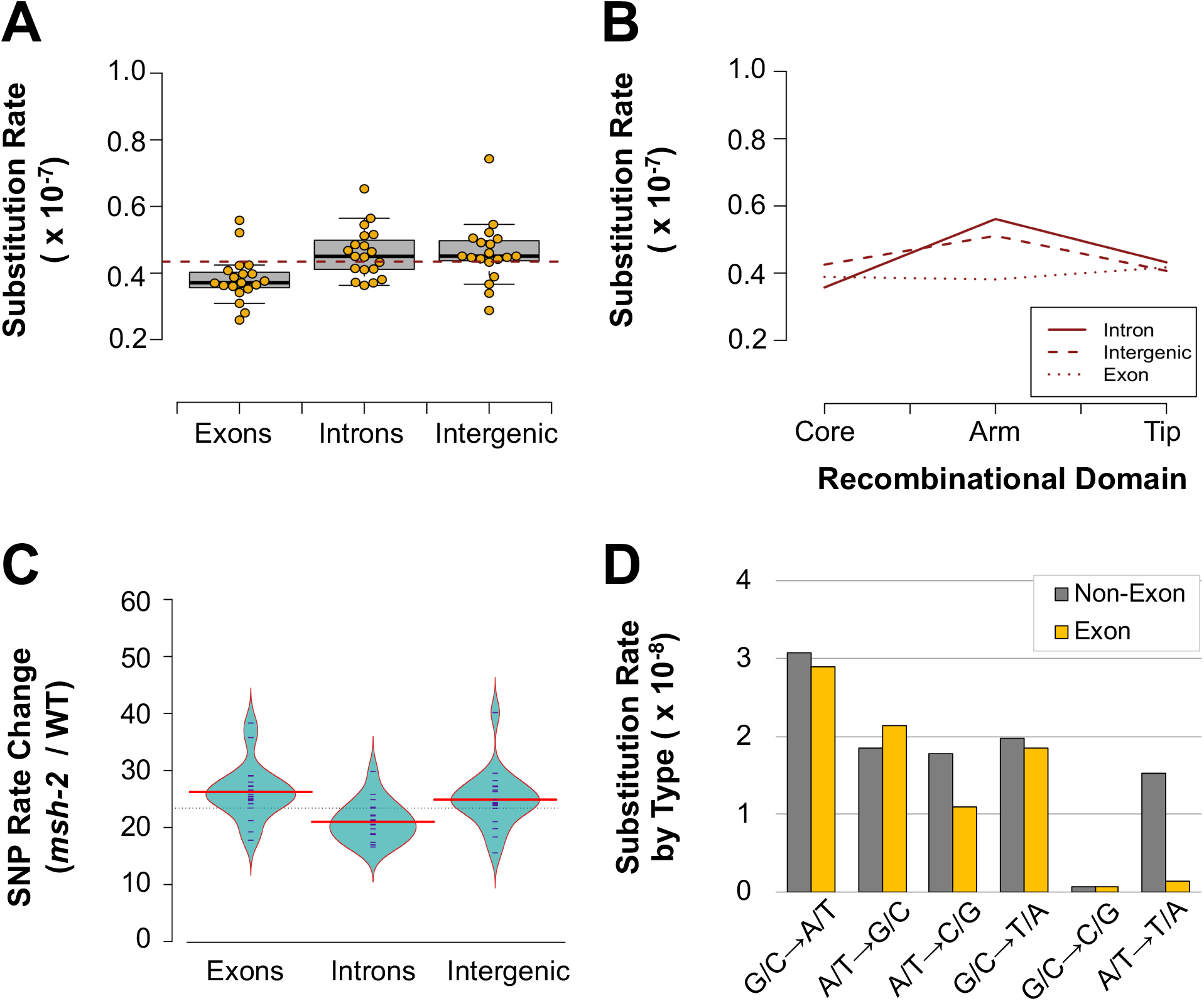
Association between coding content with the rate and spectrum of substitutions in *msh-2* knockdown MA lines. *(A)* Exons appear to be associated with lower substitution rates than noncoding DNA. However, when location in recombinational domains (cores, arms, and tips) and chromosomes are taken into account, coding content does not have a significant influence on the substitution rates. *(B)* There is a significant interaction between the coding content (exon, intron, and intergenic region) and the recombinational domains. *(C)* The increase in substitution rates in *msh-2* MA lines relative to wildtype MA lines is significantly associated with coding content. The substitution rate increase in introns is significantly less than that in exons and intergenic regions while no difference in substitution fold-changes was observed between exons and intergenic regions. *(D)* The mutational spectrum of exons is compared to that of non-exonic DNA (introns and intergenic regions). A ↔ T mutations are markedly lower in exons than the rest of the genome whereas C ↔ G mutations occur at extremely low rates throughout the genome.

The base substitution rate in chromosomal cores, arms, and tips is 3.74 (95% CI: ± 0.32), 4.77 (95% CI: ± 0.38) and 4.01 (95% CI: ± 0.59) × 10^-8^ substitutions/site/generation, respectively (**Figure 5A**). The variation between recombinational domains comprising chromosomal cores, arms, and tips is significant (3-way ANOVA: *F* = 5.40, *p* = 0.0047). However, the relative increase in base substitution rates in the *msh-2* knockdown MA lines compared to wildtype does not vary between arms, cores and tips (ANOVA: *F =* 1.67, *p =* 0.19). In contrast to our previous results for wildtype MA lines, the substitution rates during *msh-2* knockdown were significantly different between chromosomes (**Figure 5B**; 3-way ANOVA: *F =* 2.70, *p =* 0.02). In particular, the X chromosome has a lower substitution rate of 3.71 (95% CI: ± 0.51) × 10^-8^ relative to the autosomal rate of 4.33 (95% CI: ± 0.28) × 10^-8^ substitutions/site/generation (paired *t*-test: *t =* 3.28, *p =* 0.004). Lower substitution rates on the X chromosome relative to the autosomes were detected both in chromosomal cores and arms (**Figure 5C**) (2-way ANOVA: *F* = 12.53, *p* = 7.05 × 10^-4^).

**Figure 5.**
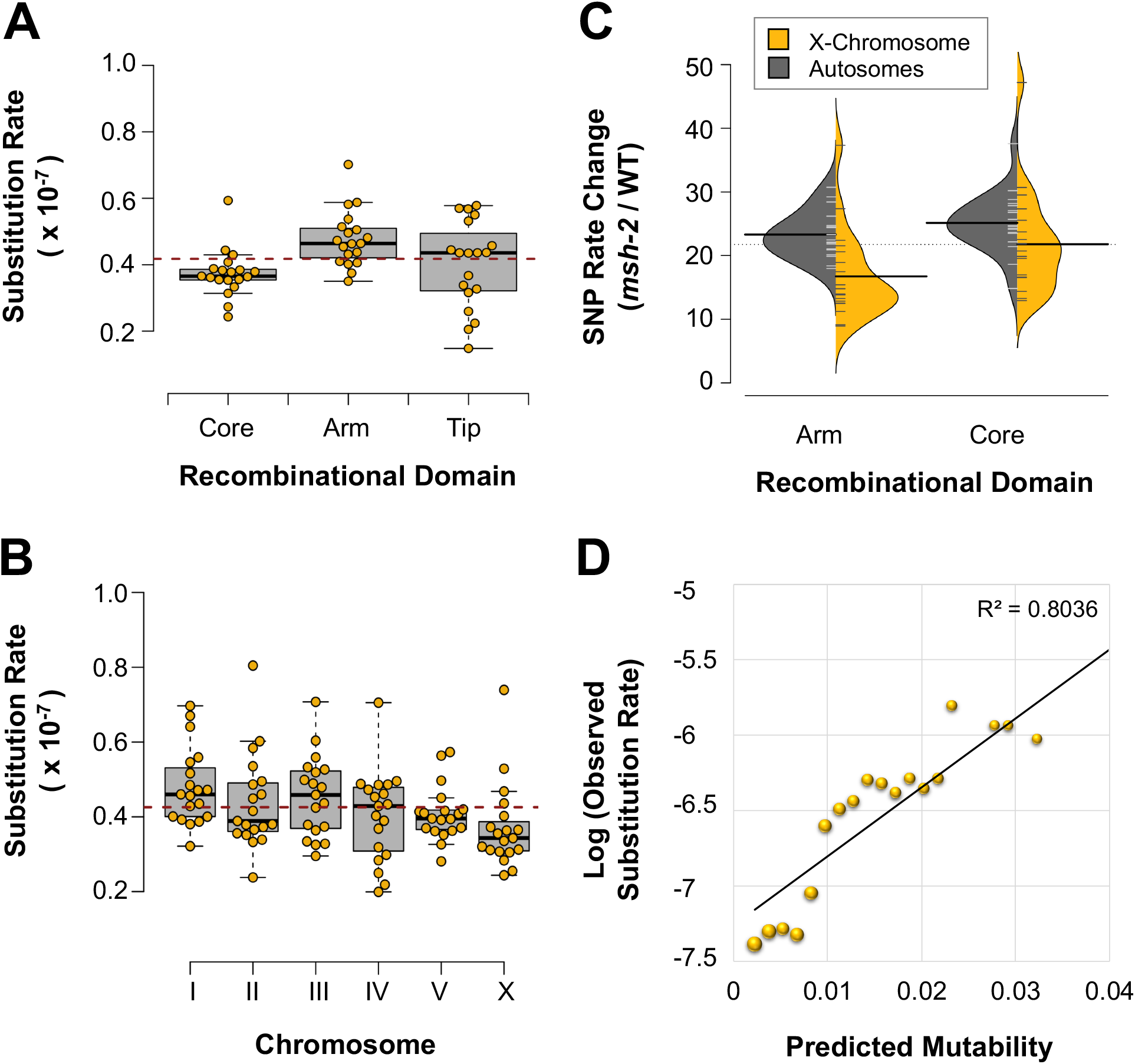
Association between genomic location and substitution rate in *msh-2* knockdown MA lines. *(A)* Significant correlation between recombinational domain (arms, cores, and tips) and substitution rates. *(B)* Significant variation in substitution rates among chromosomes. Furthermore, autosomes have higher substitution rates than the X chromosome. *(C)* The increase in substitution rates in the *msh-2* knockdown MA lines relative to wildtype MA lines is greater for the autosomes than the X chromosome, and was detected in both the chromosomal arms and cores. The distribution of the increase in substitution rates for the autosomes and X chromosome is shown in gray and yellow, respectively. *(D)* Three variables related to sequence context (repeat sequence, sequence complexity, and local G+C-content flanking a site) explain 80% of site mutability.

### Predictors of mutability

Most genomic features (chromosomal location, coding state, recombination domain, rate, and germline expression) were poor predictors of mutability in the *msh-2* MA lines using a regularized logistic regression approach (**Supplemental Figure S4A**; **Supplemental Table S2**). The presence of sequence repeats (Odds Ratio, OR = 4.42) was the best predictor for mutability (**Supplemental Figure S4A**; Konrad et al. 2019), as were certain nucleotide triplets surrounding the focal site. Local G+C-content negatively affected the mutability in the *msh-2* MA lines (OR = 0.88). However, this effect was much less than that observed in the wildtype MA lines (**Supplemental Figure S4A**). Nonetheless, the strongest positive effects of nucleotide triplets on the substitution rate in the *msh-2* lines were detected when C and G focal bases were flanked by C and G nucleotides (5′−GCC−3′/5′− GGC−3′ and 5′−CGC−3′/5′−GCG−3′; OR = 4.90 and 4.11, respectively). The relationship between several nucleotide triplets and mutability differed between the *msh-2* and wildtype MA lines, with some shifting from positive effects to negative effects and vice versa (**Supplemental Figure S4A**). With the exception of 5′−AAT−3′/5′−ATT−3′ (OR = 2.22), all other A+T triplets had negligible or negative effects on the mutability of a site (**Supplemental Table S2**; **Supplemental Figure S4A**). Most triplets containing at least two-thirds C+G bases increased mutability, while most triplets with two-thirds A+T bases reduced mutability.

An average mutability of 0.04 encompassed 100% of all SNPs and 99.9% of all genomic sites (**Supplemental Figure S4B**). The model can account for 78.8% of the variance in mutability using the same predictors as previously employed for wildtype MA lines (**Supplemental Figure S4C**; Konrad et al. 2019). However, 80.4% of the mutational variance could be accounted for if only a subset of three predictors related to sequence context (sequence repeat, G+C-content, and sequence complexity) were used to predict mutability (**Figure 5D**).

### Variation in indel rates across the genome

The indel rates observed in the *msh-2* knockdown MA lines were dependent on genomic location (**Figure 6A**-**C**). Exons had lower indel rates than introns and intergenic regions (**Figure 6A**; 3-way ANOVA: *F* = 886.82, *p* < 2.0 × 10^-16^). Additionally, rate changes due to *msh-2* knockdown relative to the wildtype lines differed significantly between exons and noncoding DNA (**Figure 6B**; ANOVA: *F =* 135.8, *p* < 2 × 10^-16^). The increase in indel rates was substantially greater in introns and intergenic regions relative to exons (Tukey’s Multiple Comparisons: introns versus exons, *p =* 0.00; intergenic regions versus exons regions, *p =* 0.00). These differences between exons and noncoding DNA are likely due to differential sequence complexity and composition in exons relative to introns and intergenic regions, as exons have higher G+C-content and fewer homopolymeric A/T runs.

**Figure 6.**
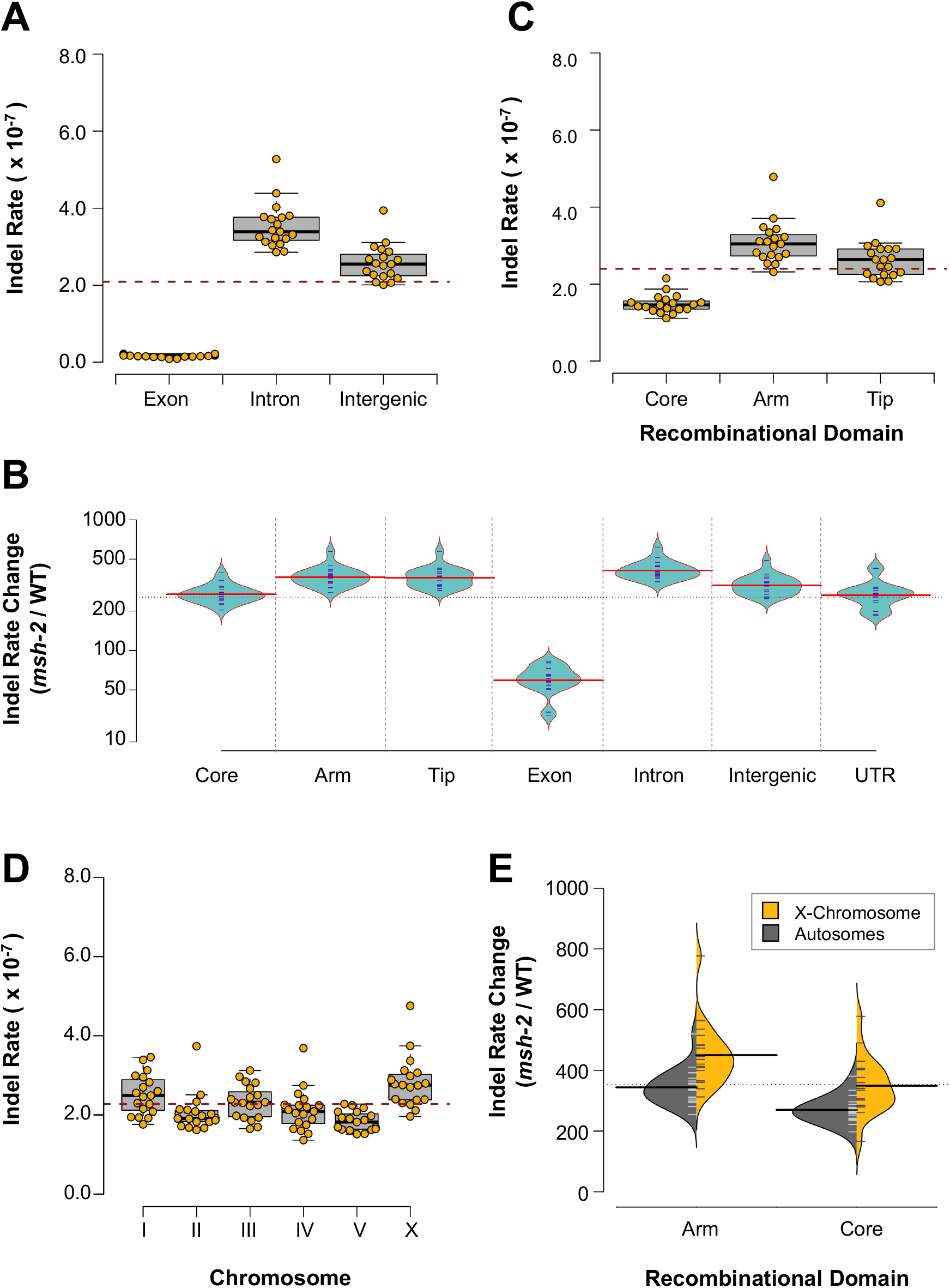
Association between genomic location and small indel rate in *msh-2* knockdown MA lines. *(A)* Indel rates differ significantly with coding content. *(B)* The relative increase in indel rates in *msh-2* MA lines relative to wildtype MA lines varies significantly between recombinational domains and coding content. While the relative increase in indel rate is significantly greater in arms and tips than in cores, the relative increase is not significantly different between arms and tips. The relative increase in the indel rate in the *msh-2* MA lines is significantly less in exons than noncoding DNA. *(C)* Indel rates differ significantly with recombinational domain. *(D)* Indel rates varied significantly by chromosome, and between autosomes and the X chromosome. *(E)* The relative increase in the indel rate in the *msh-2* knockdown MA lines relative to the wildtype MA lines was significantly greater in the X chromosome than in the autosomes and was observed on both arms and cores.

Similarly, chromosomal cores had lower indel rates than arms and tips (**Figure 6C**; 3-way ANOVA: *F* = 108.67, *p* < 2.0 × 10^-16^). Furthermore, the relative increase in indel rates during *msh-2* knockdown relative to the wildtype MA lines was greater in the arms and tips relative to the cores (**Figure 6B**; ANOVA: *F =* 15.75, *p* < 4.09 × 10^-6^; Tukey’s Multiple Comparisons: *p =* 0.00). The distribution of indels across the recombinational regions of the chromosome was not significantly different between deletions and insertions (Fisher’s Exact Test: *p =* 0.47). As discussed above for exons, gene dense cores harbor a lower proportion of mutation-prone A/T homopolymeric runs than do arms, and hence the indel rates in cores did not increase as much as in arms.

The distribution of indels showed significant variation across chromosomes (**Figure 6D**; 3-way ANOVA: *F =* 4.22, *p =* 8.41 × 10^-4^). In contrast to nucleotide substitutions, the X chromosome appears to have a higher indel rate than the autosomes (*t*-test: *t =* 4.06, *p =* 3.68 × 10^-4^). Furthermore, the increase in the indel rate on the X chromosome was greater than that on the autosomes in the *msh-2* lines relative to the wildtype MA lines, and this difference was consistent across both cores and arms (**Figure 6E**; 2-way ANOVA: *F* = 26.55, *p* = 2.16 × 10^-6^).

### Deletions outnumber insertions and a preponderance of single base indels

*msh-2* knockdown MA lines have significantly elevated indel rates relative to wildtype MA lines (**Figure 1B**; *t*-test: *t =* 26.10, *p =* 9.28 × 10^-16^), but the pattern of relative increase differs between insertions and deletions. The average deletion rate of 1.47 (95% CI: ± 0.13) × 10^-7^/site/generation is significantly higher than the average insertion rate of 7.59 (95% CI: ± 0.50) × 10^-8^/site/generation in the *msh-2* knockdown MA lines (**Supplemental Figure S5A**; *t*-test: *t =* 10.07, *p =* 5.75 × 10^-10^). While the wildtype MA lines contained an average of 3.75 deletions for each insertion, *msh-2* knockdown resulted in a significantly lower ratio of 1.94 (**Supplemental Figure S5B**; *t*-test: *t =* 2.61, *p =* 0.019). The average increase in the insertion rate during *msh-2* knockdown compared to wildtype lines is significantly higher (422×) than that for deletions (288×) (**Supplemental Figure S5C**; *t*-test: *t =* 6.89, *p* = 4.84 × 10^-8^). In the *msh-2* MA lines, the shortest indels (1-2 bp in length) exhibited the greatest increase in mutation rate relative to wildtype MA lines (3-way ANOVA: *p* < 2.0 × 10^-16^ for type, size, and the interaction of the two) and indel rate fold changes between *msh-2* and the wildtype (**Supplemental Figure S5D**; 3-way ANOVA: *p* < 2.0 × 10^-16^ for size, *p* = 2.84 × 10^-12^ type, and *p* = 1.03 × 10^-10^ for the interaction between size and type). While the frequency of ≥ 3 bp indels are increased 26×, 1-2 bp indels are increased 446× (**Supplemental Figure S5D**).

**Supplemental Figure S6** compares the size distributions of small indels in the *msh-2* knockdown and wildtype MA lines (Konrad et al. 2019). There was no difference in the average size of insertions (1.06 bp) and deletions (1.08 bp) in the *msh-2* knockdown lines (*t*-test: *t =* 1.32, *p =* 0.19). However, the overall indel size distribution in the *msh-2* knockdown MA lines had a far greater proportion of 1-bp indels relative to the wildtype MA lines (**Supplemental Figures S6A** and **B**; *t*-test: *t =* 131.9, *p* < 2.2 × 10^-16^). Single nucleotide indels comprise 97% (6960/7180) of all deletions and 95% (3513/3681) of all insertions, yielding a net loss of 3,447 bp. Indels >1-bp yield a further net loss of 381 bp. Across all 19 *msh-2* knockdown lines, this deletion bias amounts to a net loss of 3,828 bp, an average of 201.5 bp per genome, and 5-bp per genome per generation. This net loss is ∼21× higher than the wildtype rate of 0.24 bp per genome per generation (Konrad et al. 2019).

### G/C homopolymeric runs are more prone to indel mutations than A/T runs

G/C homopolymeric runs incur, on average, more indels per bp than A/T homopolymeric runs (**Supplemental Figure S7A**). The frequency of homopolymeric A/T runs in the genome is significantly higher than those of G/C nucleotides (**Supplemental Figure S7B**), which explains the higher occurrence of A/T indels despite the greater propensity of G/C homopolymeric runs to gain or lose a base pair. Both, indel type (deletions *vs.* insertions: 3-way ANOVA: *F* = 47.45, *p* = 1.77 × 10^-9^) and base pair type (A/T *vs.* G/C: 3-way ANOVA: *F* = 13.22, *p* = 5.16 × 10^-4^) have a significant effect on the per base pair indel rates. G/C homopolymeric runs have significantly higher rates of deletions than A/T homopolymeric runs (Tukey’s Multiple Comparisons: *p* = 0.009), while insertions do not differ significantly (Tukey’s Multiple Comparisons: *p* = 0.24). Additionally, neither A/T nor G/C insertion and deletion rates are constant across homopolymeric runs of different lengths (**Supplemental Figures S7C and S7D**). Shorter G/C runs display higher insertion rates whereas longer G/C runs exhibit higher deletion rates. Additionally, deletion rates increase with homopolymeric run lengths up to approximately 10 bp (G/C runs) and 11 bp (A/T runs). This heterogeneity in insertion and deletion rates in homopolymeric G/C runs results in a net gain of bases in runs < 9 bp, and a net loss of bases in runs > 9 bp (**Supplemental Figure S7D**).

### Dinucleotide microsatellites differ in indel dynamics from homopolymeric runs

Dinucleotide and polynucleotide repeats have, on average, slightly more insertions per nucleotide run than deletions (**Supplemental Figure S7E**). However, heterogeneity in the relative deletion and insertion rates between individual dinucleotide run types is apparent once the different dinucleotide runs are compared to one another (**Supplemental Figure S7F**). AC and AG dinucleotide runs experience slightly higher deletion rates than insertion rates, and CG/GC runs contained only deletions. In contrast, the insertion rate in AT/TA microsatellites was more than two-fold higher than the deletion rate.

### Mitochondrial mutations during msh-2 knockdown

Five mitochondrial mutations were detected across the 19 *msh-2* knockdown MA lines. All five mutations are heteroplasmic, although their respective intracellular frequencies ranged from relatively low (3.4 to 4.9%) to nearing fixation (81.7 to 93.7%) (**Table 3**). A nonsynonymous substitution in the *ND4L* gene resulting in a leucine to proline replacement had reached an approximately 94% frequency in MA line 16. Additionally, a deletion spanning 1,034 bp which removed the 3′ end of *COIII*, entire *tRNA-Thr* and the 5′ end of *ND4*, reached a frequency of ∼82% in MA line 16. Two frameshift mutations in the *ND5* gene occurred independently in two MA lines (∼5% and ∼4% frequency in MA line 4 and 34, respectively). Both of these frameshift mutations are single nucleotide insertions in the same homopolymeric run which we previously identified as a mutational hotspot for small indels in the *C. elegans* mitochondrial genome (Konrad et al. 2017). The number and intracellular frequencies of these mitochondrial mutations yield an overall mitochondrial mutation rate of 2.18 10^−7^ /site/generation (95% CI: ± 4.04 × 10^−7^) (**Table 1**). Although this estimated rate in the *msh-2* knockdown MA lines is two-fold greater than that calculated for wildtype MA lines (1.05 10^−7^ /site/generation; Konrad et al. 2017), a meaningful statistical test between the two experiments is precluded owing to the presence of a large number of *msh-2* MA lines with no mitochondrial mutations (**Supplemental Figure S8A**). *msh-2 knockdown does not affect the rates and length distributions of copy-number changes*

**Table 3.** List and details of five mtDNA variants identified in the *msh-2* knockdown MA lines. Five mutations were found in four lines, while the remaining 15 lines experienced no mutation. Only two of the five mutations approached fixation, while the remaining three were detected at low heteroplasmic frequencies.

| MA | MA |  |  |  |  |  |  |
| --- | --- | --- | --- | --- | --- | --- | --- |
| Line | Generation | Position | Mutation | Effect | Gene(s) | Frequency | Context |
| 16 | 33 | 721 | T→C | Leu→Pro | *ND4L* | 0.937 | TCAAGAATCC[T]GGGTATGGTA |
| 16 | 33 | 6361-7394 | 1034 bp del<sup>a</sup> | Frameshift | *COIII, tRNA-Thr, ND4* | 0.817 | ATCATCTGGG[GTT...ATA]CATCTGGGAG |
| 38 | 43 | 8872 | C→T | Pro→Leu | *COI* | 0.034 | GTATTTAATC[C]ACTTTTATTG |
| 34 | 43 | 11722 | (T)<sub>8</sub> →(T)<sub>9</sub> | Frameshift | *ND5* | 0.040 | ATTGGATTTG[T]<sub>8</sub>ATAGGTGGAA |
| 4 | 44 | 11722 | (T)<sub>8</sub> →(T)<sub>9</sub> | Frameshift | *ND5* | 0.049 | ATTGGATTTG[T]<sub>8</sub>ATAGGTGGAA |

NOTE.— The mutations are displayed as changes on the major strand.
<sup>a</sup> del = deletion

Fifteen independent copy-number changes were detected across the 19 *msh-2* knockdown MA lines (**Table 4**). A total of 16 *partial* and *complete* protein-coding genes were duplicated, yielding an overall gene duplication rate of 1.07 × 10^-6^ per gene per generation (95% CI: ± 8.36 × 10^-6^; **Table 1**). Six *partial* and *complete* protein-coding genes were deleted, yielding an overall gene deletion rate of 3.95 × 10^-7^ per gene per generation (95% CI: ± 3.71 × 10^-7^; **Table 1**). The gene duplication and deletion rates in the *msh-2* knockdown lines are not significantly different from their counterparts in the wildtype MA lines (Konrad et al. 2018) (duplications: *t*-test: *t =* 1.01, *p =* 0.33; deletions: *t*-test: *t =* 1.18, *p =* 0.26) (**Table 1**). One of the copy-number variants (CNV) was a complex event comprising a coupled duplication and deletion event. Four of the 15 CNVs were detected on Chromosome V, which is consistent with previous work identifying chromosome V as the most CNV-prone chromosome in *C. elegans* (Maydan et al. 2007, 2010; Thompson et al. 2013; Konrad et al. 2018). Although the duplication and deletion rates calculated for the *msh-2* knockdown MA lines appear to be somewhat lower than in wildtype MA lines, we did not test for significant differences between the two experiments due to the small number of *msh-2* MA knockdown lines that contained copy-number changes. Only eight and four of the 19 *msh-2* MA lines contained gene duplications and deletions, respectively.

**Table 4.**
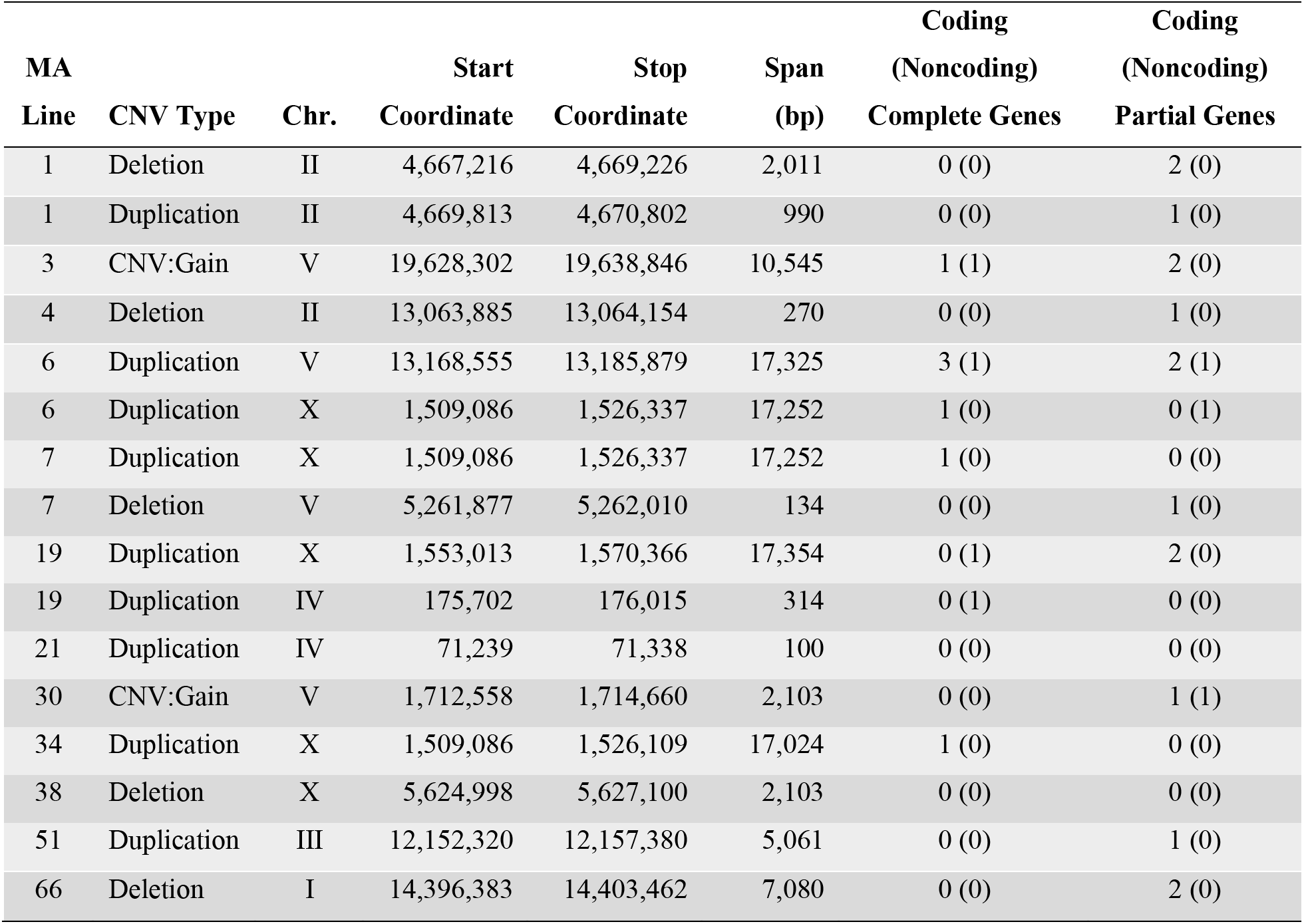
Summary of copy-number variants (CNVs) detected in the *msh-2* knockdown MA lines. The majority of copy-number changes are either novel duplications or copy-number gains in preexisting multicopy regions. Only five deletions were detected, with smaller spans than the duplications. Consequently, not a single complete gene was deleted, while multiple complete genes were duplicated throughout the MA phase.

| MA |  |  | Start | Stop | Span | Coding<br>(Noncoding) | Coding<br>(Noncoding) |
| --- | --- | --- | --- | --- | --- | --- | --- |
| Line | CNV Type | Chr. | Coordinate | Coordinate | (bp) | Complete Genes | Partial Genes |
| 1 | Deletion | II | 4,667,216 | 4,669,226 | 2,011 | 0 (0) | 2 (0) |
| 1 | Duplication | II | 4,669,813 | 4,670,802 | 990 | 0 (0) | 1 (0) |
| 3 | CNV:Gain | V | 19,628,302 | 19,638,846 | 10,545 | 1 (1) | 2 (0) |
| 4 | Deletion | II | 13,063,885 | 13,064,154 | 270 | 0 (0) | 1 (0) |
| 6 | Duplication | V | 13,168,555 | 13,185,879 | 17,325 | 3 (1) | 2 (1) |
| 6 | Duplication | X | 1,509,086 | 1,526,337 | 17,252 | 1 (0) | 0 (1) |
| 7 | Duplication | X | 1,509,086 | 1,526,337 | 17,252 | 1 (0) | 0 (0) |
| 7 | Deletion | V | 5,261,877 | 5,262,010 | 134 | 0 (0) | 1 (0) |
| 19 | Duplication | X | 1,553,013 | 1,570,366 | 17,354 | 0 (1) | 2 (0) |
| 19 | Duplication | IV | 175,702 | 176,015 | 314 | 0 (1) | 0 (0) |
| 21 | Duplication | IV | 71,239 | 71,338 | 100 | 0 (0) | 0 (0) |
| 30 | CNV:Gain | V | 1,712,558 | 1,714,660 | 2,103 | 0 (0) | 1 (1) |
| 34 | Duplication | X | 1,509,086 | 1,526,109 | 17,024 | 1 (0) | 0 (0) |
| 38 | Deletion | X | 5,624,998 | 5,627,100 | 2,103 | 0 (0) | 0 (0) |
| 51 | Duplication | III | 12,152,320 | 12,157,380 | 5,061 | 0 (0) | 1 (0) |
| 66 | Deletion | I | 14,396,383 | 14,403,462 | 7,080 | 0 (0) | 2 (0) |

### rDNA copy-number experiences more random variation in msh-2 knockdown lines

Ribosomal RNA genes (18s, 28s and 5.8s) are encoded in long tandem arrays at the end of Chromosome I in *C. elegans* (*C. elegans* Sequencing Consortium 1998). This region has previously been shown to exhibit extensive copy-number variation between nematodes, natural *C. elegans* isolates, and experimental MA lines (Bik et al. 2013; Thompson et al. 2013; Konrad et al. 2018). There is considerable variation in the estimated number of rRNA genes between the *msh-2* knockdown MA lines, ranging from ∼72 to 207 copies. The *msh-2* MA lines harbor significantly higher rDNA copy-number than the wildtype MA lines (**Supplemental Figure S8B**; *t =* 3.26, *p =* 0.0025). However, the variation in rDNA copy-number after the conclusion of each experiment did not differ significantly between the *msh-2* knockdown and wildtype MA lines (*F* = 1.04, *p* = 0.99). The wildtype MA lines were propagated for up to 409 generations, with an average of 361 generations per line, whereas the *msh-2* MA lines were maintained for up to 50 generations with an average of 40.3 generations per line. When the number of generations in each experiment is taken into account, there is significantly greater divergence of rDNA copy-number per generation in the *msh-2* lines MA lines relative to wildtype MA lines (**Supplemental Figure S8C;** *F* = 70.6, *p* = 1.34 × 10^-11^). This difference could be the result of higher recombination rates within the rDNA gene arrays of the *msh-2* knockdown MA lines.

Alternatively, the difference could be the result of higher ancestral rDNA copy-number in *msh-2* knockdown MA lines compared to their wildtype counterparts, and hence a higher probability or opportunity for unequal recombination to occur. The average copy-number change relative to the ancestral state was significantly different between the wildtype and *msh-2* knockdown lines (*t*-test: *t =* 2.57, *p =* 0.015). The *msh-2* MA lines had an average loss of seven copies of rDNA per line, which was not significantly different from a zero net loss (**Supplemental Figure S8C**; *t*-test: *t =* 0.95, *p =* 0.35). This stands in contrast to previous results from wildtype MA lines which had an average increase of 20 rDNA copies per line (Konrad et al. 2018) (**Supplemental Figure S8B**; *t*-test: *t =* 2.62, *p =* 0.02).

## DISCUSSION

DNA repair systems contribute to the evolution of genomes in multiple ways. The reduction in mutation rate by DNA repair limits genome degradation and yet, at the same time, can limit the supply of potentially beneficial mutations. The specificity of repair systems contributes to the evolution of base composition as well as the prevalence and distribution of genomic features such as the length and types of DNA sequence repeats. The efficiency and specificity of DNA repair can be constrained by the chemical and spatial configuration of mismatches in the double helix as some mismatches are more easily recognized than others. However, DNA repair could also adapt to the types and frequency of mutations in the unrepaired DNA. For instance, repair systems could adapt to preferentially repair mismatches that are particularly frequent in a genome at the expense of mismatches that are rare, or they could adapt to repair mutations that are more likely to be harmful, such as frameshifts or transversions. The specificity of DNA repair in a species would then be a function of the tradeoffs between structural constraints, the fitness cost of spontaneous mutations in the absence of repair and the cost of repair itself. Finally, the effective population size of species can set limits to adaptation of DNA repair systems. The analyses of mutation rates in DNA repair-deficient organisms increases our understanding of how spontaneous mutations arise and helps identifying DNA sequence features that are more or less susceptible to mutations. Furthermore, a comparative analysis of the mutation spectrum in repair-proficient and -deficient organisms contributes to our understanding of the DNA repair systems themselves and their evolution.

This study provides an evolutionary genome-wide view of the effect of mismatch repair impairment on the mutational landscape in *C. elegans* by severely limiting the assembly of the *mutSα* complex through RNA-interference of the *msh-2* gene in conjunction with an MA experimental design. This combination of (i) the near elimination of natural selection on spontaneous mutations, and (ii) knockdown of a key mismatch repair gene permits a more refined understanding of the raw rate and spectrum of mutational input prior to mismatch repair. We compared the mutation rates and spectra in this set of MMR-impaired obligately-outcrossing *C. elegans* MA lines to those observed in another spontaneous MA experiment with the wildtype N2 selfing strain of *C. elegans* (Konrad et al. 2017, 2018, 2019). As far as we know, there is no evidence to suggest that mutation rates and spectra differ between self-fertilizing *C. elegans* and the outcrossing populations employed in this study.

### *msh-2* knockdown is effective and increases the genome-wide nuclear mutation rate by more than 100-fold

RNAi-induced *msh-2* knockdown in *C. elegans* MA lines comprising this study resulted in a nuclear mutation rate of 2.65 × 10^-7^ /site/generation (SNPs and small indels), representing a 105× increase over the comparable mutation rate of 0.0252 × 10^-7^ /site/generation in the *C. elegans* wildtype, MMR-proficient genetic background (Konrad et al. 2019). The rates of synonymous and nonsynonymous mutations were not significantly different from each other in our MA lines, which is consistent with negligible purifying selection. A comparison of prior analyses of MA lines using Sanger sequencing of ∼20 kb (0.02%) of the *C. elegans* genome suggested a ∼48× increase in mutation rates in *msh-2* and *msh-6* knockout lines of *C. elegans* (Denver et al. 2005, 2006). Denver et al. (2006) additionally conducted a partial genomic study of two additional excision repair pathways (BER and NER) in *C. elegans* which exhibited a 17−28× increase in overall mutation rates, suggesting that the MMR pathway plays the lead role in minimizing the mutation load in *C. elegans*.

The nuclear base substitution rate of 4.22 × 10^-8^ /site/generation of the *msh-2* knockdown MA lines was ∼23× greater than that observed for wildtype MA lines by Konrad et al. (2019). As is the case for *mutS* heterodimers, only one *mutL* complex, MutLα, encoded by two *mutL* homologs *mlh-1* and *pms-1*, is present in *C. elegans*. Hence, defects in MutLα function are expected to lead to similar mutational spectra as those in the MutSα complex. However, *mlh-1*Δ and *pms-1*Δ knockout mutants exhibited substantially greater increase in the base substitution rate (∼75×; Meier et al. 2018) compared to the ∼23× increase observed by us under *msh-2* knockdown. The most striking difference in mutation rate between the *C. elegans msh-2* knockdown MA lines in this study relative to wildtype MA lines (Konrad et al. 2019) was the ∼328× increase in small indels (**Figure 1**). The knockout of *mutL* homologs *mlh-*1 and *pms-1* in *elegans* each resulted in ∼440× increase in the small indel rate (Meier et al. 2018). One possibility for the differential increase in base substitution and indel rates under *mutL-* (Meier et al. 2018) versus *mutS*-impairment (this study) is that defects in these different components of MMR have different consequences for mutation and repair. Alternatively, RNAi-induced knockdown in our study did not fully inactivate *msh-2*. However, differences in mutation rate between *mutS and mutL* lines in the same species have been noted before (Lee et al. 2012; Long et al. 2016).

### Interspecific variation in the efficiency and specificity of MMR systems

Genome-wide mutations rates (base substitutions and indels) in an MMR-deficient background can be quite variable across as well as within species, reflecting both species-specific differences in the efficiency and specificity of mismatch repair and the influence of genetic background within a species, in addition to methodological differences in the analysis of mutations (Long et al. 2018, **Supplemental Tables S3; Supplemental Figures S9A-D**). MMR-deficiency increases the genome-wide mutation rate by one to three orders of magnitude. We further compare mutation rates under *msh-2* impairment or deficiency in an MA setting in two other eukaryotes namely *S. cerevisiae* and *A. thaliana.* The increase in base substitution rates under *msh-2* deficiency in *C. elegans* (23×; this study) is more similar to *S. cerevisiae* (15-27×; Lang et al. 2013; Lujan et al. 2014; Serero et al. 2014) than to the other multicellular eukaryote *A. thaliana* (170×; Belfield et al. 2018). The relative increase in small indels under *msh-2* deficiency in *C. elegans* (328×; this study) is much lower relative to *S. cerevisiae* (703-3,500×; Lang et al. 2013; Serero et al. 2014) and *A. thaliana* (1,000×; Belfield et al. 2018). Irrespective, the three eukaryotic species exhibit two to three orders of magnitude increase in small indel rates, suggesting that the MMR system in eukaryotes is far more efficient in the repair of small indels relative to base substitutions. The consequences of mismatch-repair deficiency for indels relative to base substitutions vary greatly between taxa (**Supplemental Table S3**). In the six species of prokaryotes thus far studied, MMR-defective strains for four species (*V. fischeri*, *B. subtilis*, *P. fluorescens* and *E. coli*) had ratios of fold-change increases in the small indel rate relative to the base substitution rate ranging from 0.32 to ∼1.0, revealing either (i) a base substitution-bias or (ii) approximately equal rates of indels and base substitutions under MMR-deficiency. Two prokaryotic species, *V. cholerae* and *radiodurans*, however, had ratios of increases in the small indel rate relative to the base substitution rate ranging from 1.67 −5.25 signifying a small indel bias relative to base substitutions under MMR-deficiency. *C. elegans* shares similarities with the other two eukaryotes, *A. thaliana* (ratio of ∼5; calculated from Belfield et al. 2018) and *S. cerevisiae* (ratios of ∼233 calculated from Lang et al. 2013, and ∼31 calculated from Serero et al. 2014) which also display a far greater increase in small indels relative to base substitutions in an MMR-impaired background. In eukaryotes, MMR-deficiency appears to lead to a far greater increase in the number of indels relative to base substitutions, suggesting a functional divergence in the MMR machinery of eukaryotes and prokaryotes. However, elucidating the causes of variation in the specificity of mismatch repair requires a far broader taxonomic sample of studies investigating the consequences of mismatch-repair deficiency on the mutational spectrum and fitness.

### Dependence of small indel rate on base composition and length of homopolymeric run

In our study, while the overall indel mutation rate increased ∼328×, single base pair insertion and deletion rates increased by 638× and 517×, respectively. These 1 bp indels were predominantly found in homopolymeric runs. The shortest indels (1-2 bp in length) are both (i) most common, and (ii) exhibit the greatest increase in mutation rates in our *msh-2* MA lines compared to wildtype MA (Konrad et al. 2019). The preference for repairing the shortest indels by MMR has also been observed in prokaryotes (Long et al. 2018). The average increase in the insertion rate during *msh-2* knockdown compared to wildtype lines was significantly higher (422×) than that for deletions (288×). Although both the wildtype and *msh-2* MA lines exhibited a deletion bias, it was significantly lower in the *msh-2* MA lines. These results suggest that MMR in *C. elegans* is better at detecting or repairing small insertions relative to deletions.

It is well-established that runs of identical nucleotides are mutational hotspots for small indels (Streisinger et al. 1966; Lee et al. 2012). Runs of G/C are significantly less stable than those of A/T in the *msh-2* lines, especially with respect to deletions. Higher indel rates in homopolymeric runs of G/C than A/T have been previously observed in a partial genomic analysis (∼20 kb) in *C. elegans* (Denver et al. 2005) and in *S. cerevisiae* MMR-deficient lines (Gragg et al. 2002; Lang et al. 2013). Additional differences between G/C and A/T homopolymeric runs in our *C. elegans* MMR-impaired lines include the tendency for an insertion-bias in shorter (<9 bp) G/C runs versus the tendency towards a deletion-bias in A/T runs, regardless of the length of the run. MMR-deficient lines of *E. coli* also exhibit an insertion-bias in G/C homopolymeric runs (Lee et al. 2012).

Lee et al. (2012), among others, have previously proposed that the indel rate in homopolymeric runs is dependent on the length of the run. The indel rates in homopolymeric runs in *E. coli mutL*, *Vibrio fischeri mutS* and *V. cholerae mutS*, increase with the length of a run of up to 10 bp (Lee et al. 2012; Dillon et al. 2017). However, the relationships between indel rates and the length of the homopolymeric runs in bacteria may have been limited by a small number of runs with >10 bp. In *C. elegans*, the indel rate peaks in 11 bp runs and is lower in runs that were either shorter or longer, both in wildtype MA lines (Konrad et al. 2019) and *msh-2* MA lines comprising this study. However, when the indel frequency was analyzed separately in G/C and A/T runs in wildtype and *msh-2* MA lines of *C. elegans*, the A/T runs followed this same pattern whereas the G/C indel rates peaked in runs of 8 bp. Interestingly, the indel rates in homopolymers in *A. thaliana*, which were primarily A/T, also peaked at 11 bp (Belfield et al. 2018). The fact that we see the same relationship between indel rates and the length of A/T homopolymeric runs in *C. elegans* and *A. thaliana* could be a mere coincidence. However, it merits further investigation into possible general rules of short indel dynamics and their consequences for genome evolution.

### Dinucleotide microsatellites differ in indel dynamics from homopolymeric runs

In contrast to homopolymeric repeats, dinucleotide and polynucleotide repeats had an average tendency for more insertions than deletions in our MMR-impaired MA lines. However, this tendency appeared to vary based on the specific composition of the repeat unit. For example, (AT)*_n_* dinucleotide repeats appeared to have 2× more insertions than deletions, a mutational bias that has also been observed in a genome-wide analysis of *S. cerevisiae* MMR-deficient lines (Lang et al. 2013). In contrast to (AT)*_n_* dinucleotides, (AC/GT)*_n_* and (AG/CT)*_n_* dinucleotide repeats in our MMR-deficient lines appear to have marginally higher deletion rates whereas only deletions were observed in (GC)*_n_* repeats. Lang et al. (2013) also observed a slight deletion bias, albeit nonsignificant, in (AC/GT)*_n_* repeats in yeast. Degtyareva et al. (2002) analyzed five microsatellite loci ((GT)_14_, (GT)_26_, (GT)_59_, (AAT)_28_, (AAAT)_43_) in *C. elegans msh-2* deficient lines and observed a 2.2-fold higher incidence of insertions to deletions (33 *vs.* 15) in the pooled sample. The discrepancy between our results and those of Degtyareva et al. (2002) may stem from limited sampling in the latter study as compared to a genome-wide analysis or due to different repair efficiency in knockout versus knockdown experiments.

### *msh-2* knockdown leads to increased transition bias

MMR repairs transitions more efficiently than transversions, which results in an increased transition bias in MMR-impaired MA lines. The transition to transversion ratio (Ts/Tv, henceforth) was significantly higher in the *msh-2* knockdown MA lines relative to the wildtype MA lines (Konrad et al. 2019), increasing from 0.67 to 1.12, or 1.7×. In *A. thaliana*, the Ts/Tv ratio increased 2.8× in *msh-2* MA lines (Belfield et al. 2018). The transition bias also increased in *S. cerevisiae mutS* deletions (Lang et al. 2013; Serero et al. 2014). In bacteria, the increase in the Ts/Tv ratio in MMR-deficient strains compared to wildtype ranges from 3× to 48× (Long et al. 2018, and references therein).

In MA experiments with eukaryotes, G/C → A/T transitions are the primary contributor to an increased Ts/Tv bias in MMR-deficient lines (Lang et al. 2013; Serero et al. 2014; Belfield et al. 2018). The disproportionately high share of G/C → A/T transitions of all spontaneous base substitutions have been explained by deamination of methylated cytosines (Lutsenko and Bhagwat 1999). Both *Saccharomyces* and *Arabidopsis* employ CpG methylation as a part of their epigenetic toolkit which is consistent with the differences in G/C → A/T transitions between these two species (Finnegan et al. 1996; Tang et al. 2012); *S. cerevisiae* has relatively low levels of CpG methylation (Tang et al. 2012), and exhibits a much lower contribution of G/C → A/T transitions to the mutational spectrum than *A. thaliana*, which has higher levels of CpG methylation (Takuno et al. 2016). In *C. elegans*, however, the relative contribution of G/C → A/T transitions to the substitution spectrum is much lower (25%) compared to *S. cerevisiae* (35 – 48%) or *A. thaliana* (76%) despite similar genomic G+C-content (38% in *S. cerevisiae* and 36% in *C. elegans*, and *A. thaliana*) (Lang et al. 2013; Serero et al. 2014; Belfield et al. 2018). The difference in the rates of transitions in *C. elegans* relative to other eukaryotes could be due to the lack of the CpG methylation (Simpson et al. 1986). Interestingly, *S. pombe*, which is also devoid of CpG methylation (Antequera et al. 1984) shows a similarly low contribution of G/C → A/T transitions to the mutation spectrum as *C. elegans* (Sun et al. 2016). This further implicates differences in CpG methylation in driving the variation between species in G/C → A/T transitions, and the transition bias.

The variation in the efficiency of repairing different mismatches also influences the evolution of genome base composition. In *C. elegans*, the mutation bias toward a lower GC-content is greater in the wildtype MA lines relative to their *msh-2* knockdown counterparts. A/T → G/C transitions were repaired more efficiently than G/C → A/T transitions and A/T → C/G transversions were repaired more efficiently than G/C → T/A transversions (**Figure 2B**). Hence, the MMR system contributes to the mutational bias toward a greater AT-content in *C. elegans*.

### Local sequence-context influences mutation rates in MMR-deficient lines

The DNA sequence flanking mutated bases appears to influence the probability of mutation in the *msh-2* MA lines. Both A/T and G/C base pairs are most prone to mutations when they are flanked by a 5′-G and a 3′-C. Several empirical studies of the local sequence-context on mutation rate in bacteria have found that guanines and cytosines flanking the focal nucleotide increase the probability of a mutation (Lee et al. 2012; Long et al. 2014; Sung et al. 2015; Takemoto et al. 2018). The association between flanking G+C-content and mutation rate implicates strong base pairing of flanking nucleotides in stabilizing mismatched bases (Sung et al. 2015).

Even more striking is the context-dependence of A/T → T/A transversions, which were more common when the mutated A/T base pair was flanked by a 5′-A and a 3′-T than when flanked by other bases. A/T → T/A transversions also exhibited context-dependence in the wildtype MA lines, but with a higher mutation rate when flanked by 5′-T and a 3′-A (Konrad et al. 2019; Saxena et al. 2019). In our *msh-2* knockdown MA lines, A/T → T/A transversions were particularly frequent within or adjacent to repetitive sequences, especially at the boundaries of homopolymeric runs.

### Significant variation in rDNA copy-number

In contrast to duplications and deletions of single-copy genes, the dynamics of rDNA copy-number are different between the *msh-2* knockdown and wildtype MA lines. Two independent experiments have found that rDNA copy-number increased during MA in wildtype *C. elegans* lines (Bik et al. 2013; Konrad et al. 2018). Although our *msh-2* knockdown MA lines displayed a significant variation in rDNA copy-number, there was no significant change in the average copy-number. Furthermore, there was a significant difference in the direction of change in rDNA copy-number between this study and our *C. elegans* wildtype MA experiment of Konrad et al. (2018). Could *msh-2* be involved in rDNA copy-number control and bias copy-number changes towards an increase rather than a decrease or random directional change? There are additional considerations that could potentially explain the difference between the *msh-2* knockdown and wildtype MA results. First, the ancestral rDNA copy-number differed between the two experiments. The N2 ancestor of the wildtype MA lines was estimated to have 98 copies (Konrad et al. 2018) whereas the *fog-2* ancestor of the *msh-2* knockdown MA lines contained an estimated 160 copies of rDNA. The range in rDNA copy-number may be limited by lower and upper boundaries, and because the N2 ancestor of the wildtype MA lines had fewer copies, a reduction in rDNA copy-number may more frequently fall below what is permissible without a significant reduction in fitness. Consequently, purifying selection might operate more often on rDNA copy-number losses than increases in the wildtype MA lines. This would result in the wildtype MA lines displaying evidence of copy-number increase more frequently than decrease and give the impression of a mutational bias towards a copy-number increase. Furthermore, the greater ancestral copy-number of rDNA in the *msh-2* knockdown lines may simply provide more substrate for generating new copy-number variation through recombination. This in turn would lead to more copy-number changes per generation regardless of the functionality of *msh-2*. Recently, variation in rDNA copy-number has been associated with broad changes in gene expression in *Drosophila* and humans (Paredes and Maggert 2009; Gibbons et al. 2014). Further experiments are clearly needed to understand the evolutionary dynamics, and the transcriptional and phenotypic consequence of rDNA copy-number dynamics in *C. elegans*.

### The X-chromosome has lower substitution rate and higher indel rate than the autosomes

Male-biased mutation rates have been observed in many species of mammals, bird, and plants (Whittle and Johnston 2003; Wilson-Sayres and Makova 2011; Jónsson et al. 2017). *C. elegans* has an XO sex-determination system wherein wildtype worms are predominantly self-fertilizing XX hermaphrodites with XO males in rare frequency. However, because the *msh-2* MA experiments were performed in a *fog-2* genetic background which results in obligate outcrossing and equal numbers of males and females, 2/3 of the X chromosomes are in females and 1/3 in males in each generation. This predicts a lower observed mutation rate on the X chromosome than on the autosomes if males have a higher mutation rate than females. For example, MA experiments with *Drosophila* have detected slightly higher mutation rates in males due to the X chromosome having lower mutation rate than the autosomes, although the difference was not statistically significant (Keightley et al. 2009). Interestingly, base substitution rates on the X are significantly lower than on the autosomes in the *C. elegans msh-2* MA lines, which is consistent with a male-biased mutation rate. In contrast, we found higher small indel rates on the X chromosome, consistent with a female-bias in indel rates. However, chromosome location was a poor predictor of mutation rate in the logistic regression analysis. Although the differences in mutation rate between the X chromosome and the autosomes are consistent with sex-specific differences in mutation rate, we cannot rule out the influence of chromosomal variation with respect to the number and distribution of mutable sites. In addition to differences between the X chromosome and the autosomes, other aspects of genomic location and sequence-context were associated with the mutation rate. For example, both base substitutions and indels were less likely to occur in exons than in intergenic regions, and less likely in chromosomal cores versus arms. However, as with the X chromosome to autosome comparison, recombinational domains (cores, arms and tips) and location in coding versus noncoding DNA were poor predictors of mutability. Most of the variation in mutation rate within the genome could be explained by sequence complexity, sequence repeats, and G+C-content. Furthermore, the indels were dominated by 1 bp deletions and insertions in runs of As and Ts, and the nonrandom distribution of A/T homopolymeric runs across the genome appears to explain the variation in indel rates.

In summary, this study combines a mutation accumulation experimental framework coupled with whole-genome sequencing to investigate alterations to the mutation rate and spectrum under impaired functionality of the MutS homolog, *msh-2* in *C. elegans*, at both the mitochondrial and nuclear levels. The large number of mutations, 13,986 in total, enabled the most comprehensive view of the characteristics and distribution of mutations in any given *C. elegans* genotype. Some results were surprising, such as the great rate of divergence in rRNA copy-number compared to our previous study of wildtype *C. elegans* lines, and differences in mutation rates between the X chromosome and autosomes, which is consistent with sex-specific mutation rates. These differences in *C. elegans* may be a consequence of a nonrandom distribution of mutable sites in the genome, which also affects differences between coding and non-coding DNA and between chromosome arms and cores. However, the results suggest that possible sex-specific differences in mutation rates in *Caenorhabditis* should be further investigated. The probability of mutation is influenced by several factors that are shared with other species, such as sequence complexity and flanking nucleotide G+C-content. In contrast, the sequence context of A/T to T/A transversions at the ends of homopolymeric runs have not been readily apparent in other species. MMR impairment has variable consequences in different species, which demonstrates that although the MMR system is evolutionary conserved, the specificity of MMR in different taxa can vary and is evolutionary labile. Whether the specificity of MMR is any given species is adapted to the distribution and fitness effects of endogenous DNA replication errors in those species is an open question.

## ACKNOWLEDGEMENTS

This manuscript greatly benefitted from comments provided by three anonymous reviewers. This research was supported by National Science Foundation Grant MCB-1565844 to V.K. U.B. and V.K. were additionally supported by start-up funds from the Department of Veterinary Integrative Biosciences, College of Veterinary Medicine and Biomedical Sciences at Texas A&M University. We thank Robert Waterston (University of Washington) and Donald Moerman (University of British Columbia) for their help with genome sequencing. We are grateful to Philip Green from the University of Washington for providing the program Phaster.

## Supplemental Material

**Supplemental Table S1.** Summary of *msh-2* knockdown MA lines, generations propagated under population size bottlenecks under a mutation accumulation (MA) experimental regime, and sequence coverage per genome.

| **Genotype** | **MA Line** | **MA Generations** | **Coverage** |
| --- | --- | --- | --- |
| *fog-2 / msh-2 RNAi* | 1 | 38 | 18 |
| *fog-2 / msh-2 RNAi* | 3 | 44 | 17 |
| *fog-2 / msh-2 RNAi* | 4 | 44 | 15 |
| *fog-2 / msh-2 RNAi* | 6 | 33 | 16 |
| *fog-2 / msh-2 RNAi* | 7 | 28 | 17 |
| *fog-2 / msh-2 RNAi* | 16 | 33 | 17 |
| *fog-2 / msh-2 RNAi* | 19 | 43 | 18 |
| *fog-2 / msh-2 RNAi* | 21 | 39 | 17 |
| *fog-2 / msh-2 RNAi* | 28 | 36 | 19 |
| *fog-2 / msh-2 RNAi* | 30 | 42 | 16 |
| *fog-2 / msh-2 RNAi* | 34 | 43 | 16 |
| *fog-2 / msh-2 RNAi* | 35 | 43 | 18 |
| *fog-2 / msh-2 RNAi* | 38 | 43 | 16 |
| *fog-2 / msh-2 RNAi* | 45 | 41 | 16 |
| *fog-2 / msh-2 RNAi* | 50 | 44 | 16 |
| *fog-2 / msh-2 RNAi* | 51 | 45 | 16 |
| *fog-2 / msh-2 RNAi* | 66 | 45 | 16 |
| *fog-2 / msh-2 RNAi* | 68 | 39 | 16 |
| *fog-2 / msh-2 RNAi* | 74 | 43 | 14 |
| *Ancestral fog-2 pre-MA ancestral control* | PMAC | 0 | 30 |

**Supplemental Table S2.** Fitted coefficients and Odds Ratios *(OR*) for predictor contribution to site mutability using a logistic regression approach.

| Predictor | Coefficient | Odds Ratio |
| --- | --- | --- |
| V | 0.24 | 1.27 |
| IV | 0.04 | 1.04 |
| I | 0.00 | 1.00 |
| X | 0.00 | 1.00 |
| II | -0.05 | 0.95 |
| III | -0.17 | 0.84 |
| Intron | 0.01 | 1.01 |
| 3'UTR | 0.00 | 1.00 |
| 5'UTR | 0.00 | 1.00 |
| Intergenic | -0.01 | 0.99 |
| Exon | -0.07 | 0.93 |
| Core | 0.00 | 1.00 |
| Tip | -0.01 | 0.99 |
| Arm | -0.04 | 0.96 |
| Recombination Rate | 0.03 | 1.03 |
| Germline Expressed | 0.00 | 1.00 |
| Repeat Sequence | 1.49 | 4.42 |
| Seq. Complexity (40 bps) | 0.33 | 1.39 |
| CG Content (40 bps) | -0.13 | 0.88 |
| GCC/GGC | 1.59 | 4.90 |
| CGC/GCG | 1.42 | 4.12 |
| AGC/GCT | 1.33 | 3.79 |
| AGG/CCT | 1.09 | 2.97 |
| GAC/GTC | 0.91 | 2.49 |
| AAT/ATT | 0.80 | 2.22 |
| ACG/CGT | 0.76 | 2.15 |
| CAC/GTG | 0.75 | 2.12 |
| GCA/TGC | 0.75 | 2.11 |
| AAC/GTT | 0.44 | 1.56 |
| CTC/GAG | 0.40 | 1.50 |
| CAG/CTG | 0.40 | 1.49 |
| AAG/CTT | 0.38 | 1.46 |
| ATC/GAT | 0.28 | 1.33 |
| ACC/GGT | 0.26 | 1.30 |
| ATG/CAT | 0.23 | 1.26 |
| ACT/AGT | 0.23 | 1.26 |
| CCC/GGG | 0.19 | 1.21 |
| GTA/TAC | 0.17 | 1.19 |
| ACA/TGT | 0.13 | 1.14 |
| CCG/CGG | 0.13 | 1.13 |
| GGA/TCC | 0.08 | 1.08 |
| AGA/TCT | 0.01 | 1.01 |
| CGA/TCG | -0.08 | 0.92 |
| CTA/TAG | -0.08 | 0.92 |
| TAA/TTA | -0.23 | 0.80 |
| CCA/TGG | -0.40 | 0.67 |
| GAA/TTC | -0.41 | 0.67 |
| ATA/TAT | -0.52 | 0.60 |
| TCA/TGA | -0.71 | 0.49 |
| CAA/TTG | -0.73 | 0.48 |
| AAA/TTT | -1.64 | 0.19 |

**Supplemental Table S3.**
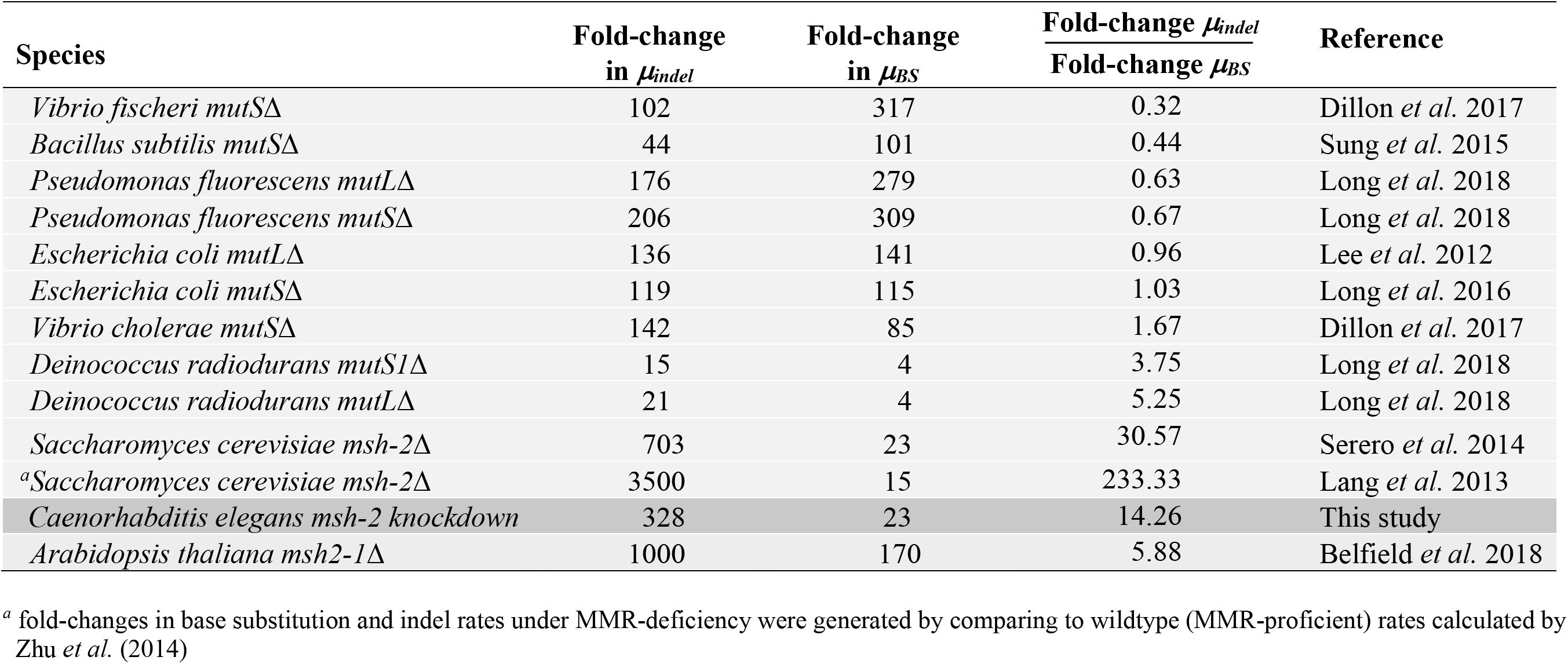
Summary of fold-change in the nuclear base substitution and small indel rates in MMR-deficient MA lines of various prokaryote and eukaryote species relative to their MMR-proficient counterparts. *μ_indel_* and *μ_BS_* refer to the mutation rates for nuclear small indels and base substitutions, respectively.

**Supplemental Table S4.** Summary of fold-change in the nuclear base substitution rates in MMR-deficient MA lines of three eukaryote species relative to their MMR-proficient counterparts.

| **Substitution type** | **Fold-increase in *C. elegans* under *msh-2 KD*** | **Fold-increase in *A. thaliana* under *msh-2 KO*** | **Fold-increase in *S. cerevisiae* under *msh-2 KO*** |
| --- | --- | --- | --- |
| G/C → A/T | 22 | 190 | 47 |
| A/T → G/C | 54 | 115 | 51 |
| G/C → C/G | 3 | 68 | 10 |
| G/C → T/A | 26 | 101 | 24 |
| A/T → C/G | 46 | 56 | 9 |
| A/T → T/A | 13 | 57 | 13 |

<sup>a</sup> fold-changes in base substitution rates under MMR-deficiency in *C. elegans* (this study) were generated by comparing to wildtype (MMR-proficient) rates calculated by Konrad *et al.* (2019)
<sup>b</sup> fold-changes in base substitution rates under MMR-deficiency in *A. thaliana* (Belfield *et al.* 2018) were generated by comparing to wildtype (MMR-proficient) rates calculated by Ossowski *et al.* (2010)
<sup>c</sup> fold-changes in base substitution rates under MMR-deficiency relative to wildtype in *S. cerevisiae* (Lujan *et al.* 2014)

**Supplemental Figure S1.**
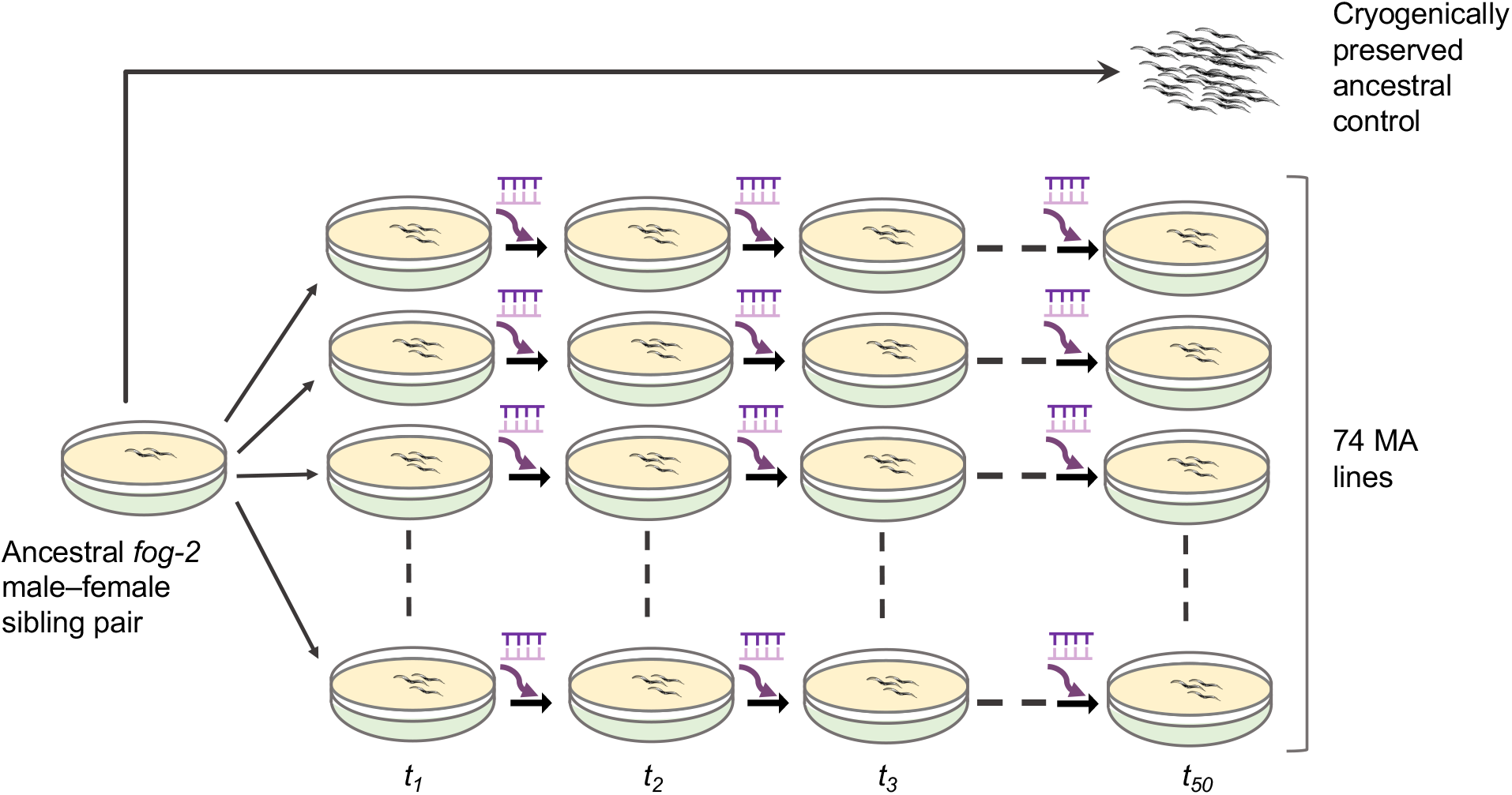
Schematic of the *msh-2* knockdown mutation accumulation (MA) experiment. The MA experiment was established from a single male-female pair of an obligatory outcrossing *fog-2* mutant strain of *C. elegans*. A trio of one virgin female and two males from the descendants of the founding pair were each used to establish 74 MA lines. The accumulation of mutations was accelerated by simultaneously (i) bottlenecking populations (*N* = 3 individuals), and (ii) RNAi-induced knockdown of MMR gene *msh-2* (symbolized by the purple RNA-RNA duplex and arrow) in each new generation. The MA experiment was conducted for up to 50 consecutive generations.

**Supplemental Figure S2.**
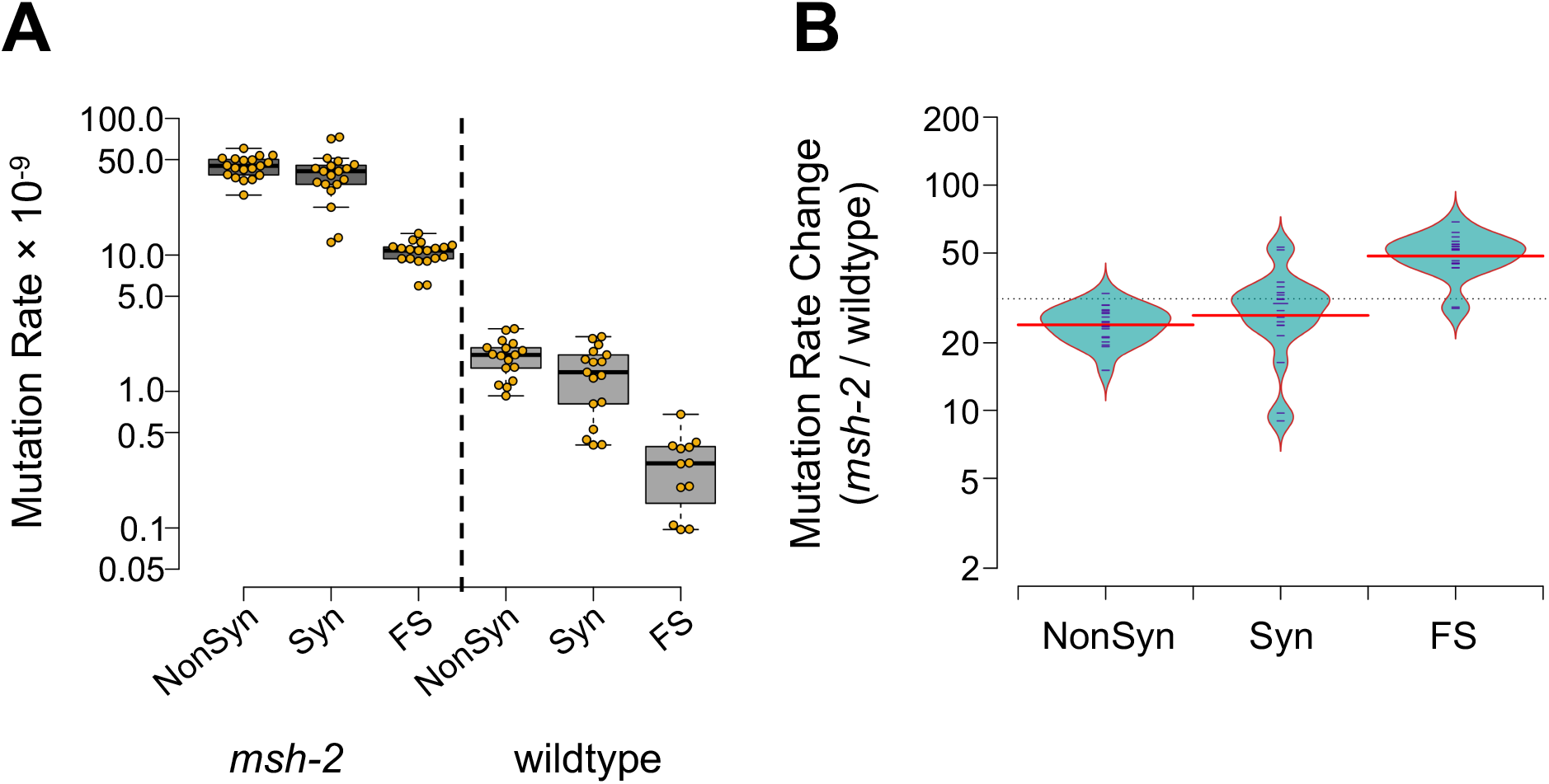
Mutation rates in exons of *msh-2* knockdown versus wild-type MA lines. (*A)* The synonymous and nonsynonymous substitution rates do not differ significantly in the *msh-2* MA lines. However, frameshift mutations in the *msh-2* MA lines were less frequent than either synonymous or nonsynonymous substitutions. The corresponding mutation rates in wildtype MA lines are shown for comparison (Konrad et al. 2019). *(B)* The frequencies of nonsynonymous, and synonymous substitutions exhibit a similar increase in *msh-2* MA lines relative to wildtype MA. The frameshift mutation rate in *msh-2* MA lines exhibits a greater increase than either nonsynonymous or synonymous rates compared to wildtype MA lines. Nonsynonymous, synonymous, and frameshift mutations are referred to as Nonsyn, Syn, and FS in the figures, respectively.

**Supplemental Figure S3.**
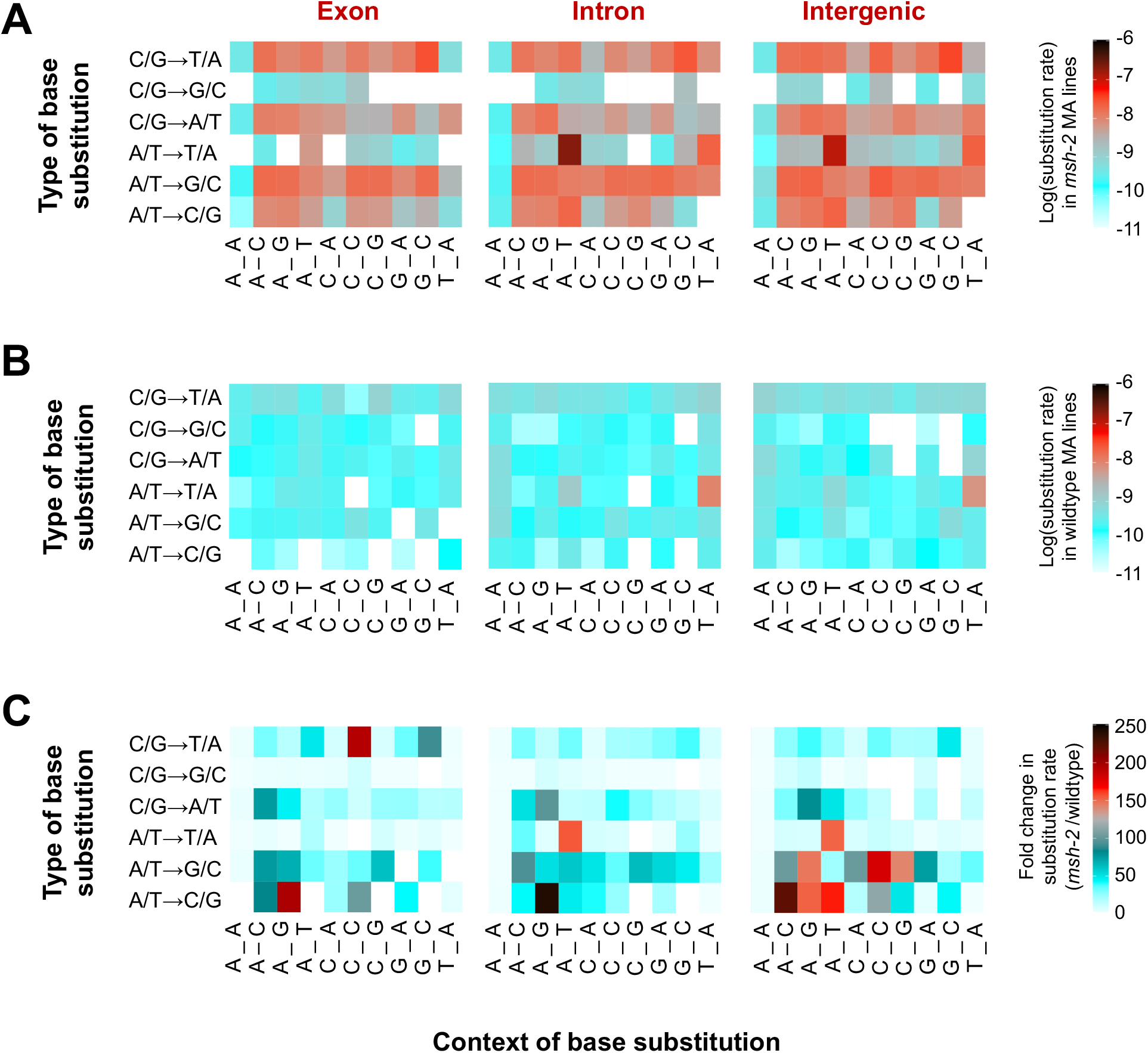
Heat maps of specific base substitution rate with their immediate flanking nucleotides. The context-dependent mutational spectra are shown for *(A) msh-2* knockdown and *(B)* wildtype MA lines. While wildtype MA lines exhibit a strong A/T → T/A mutation bias when flanked by a 5′−T and a 3′−A, *msh-2* knockdown lines exhibit an increase in the same substitution type when flanked by a 5′−A and a 3′−T. *(C)* The increase in each context-dependent substitution rate in *msh-2* versus wildtype MA lines differs between exons, introns, and intergenic regions.

**Supplemental Figure S4.**
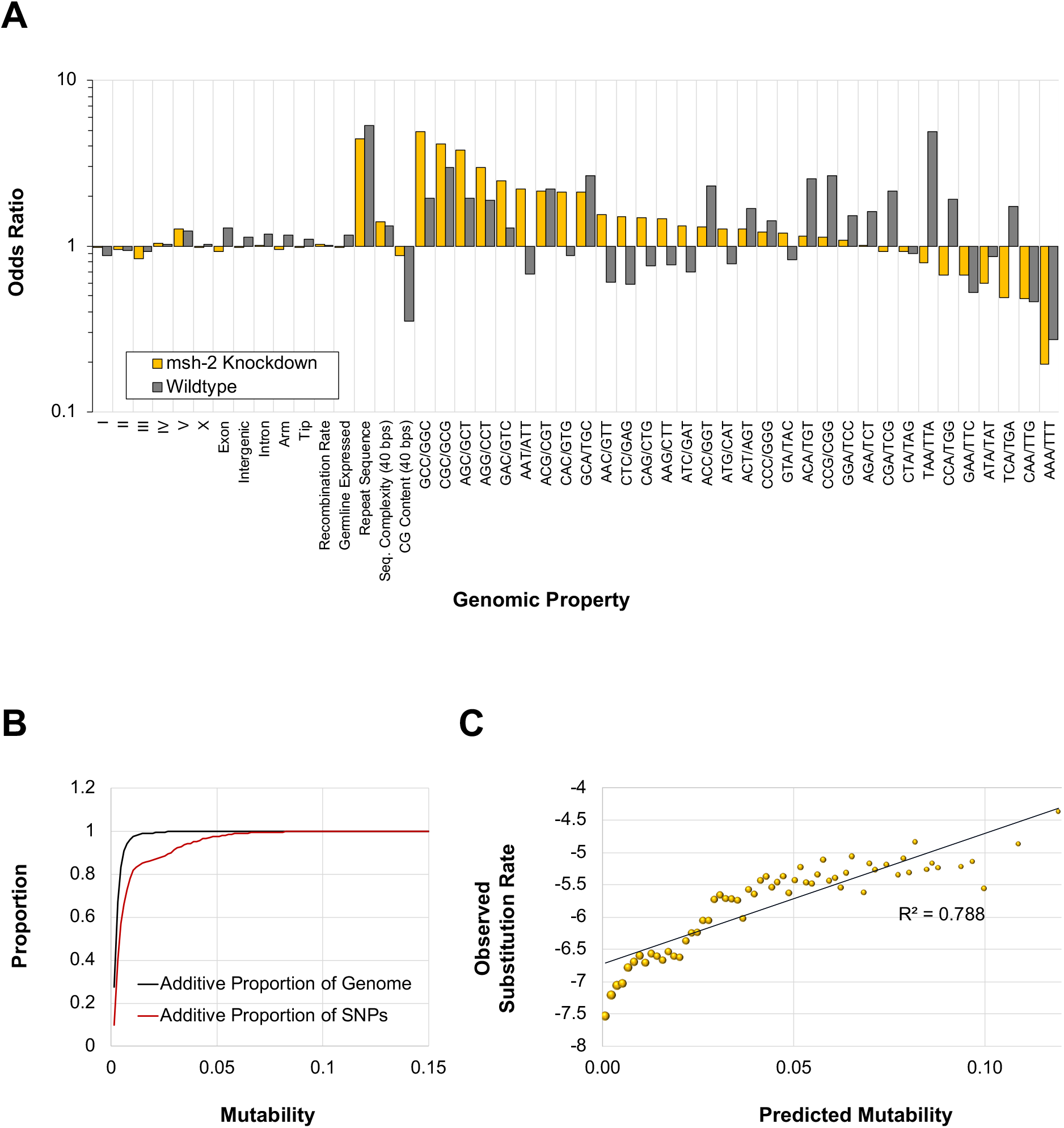
Genomic properties associated with base substitution sites. *(A)* Odds ratios of the contribution of various genomic predictors towards site mutability as estimated by a logistic regression approach. *(B)* The cumulative proportion of the genome (black) and SNPs (red) covered by increasing predicted mutabilities. *(C)* The correlation between predicted mutability and observed mutation rate is significant (*R^2^* = 0.79, *p* < 2.2 × 10^-16^).

**Supplemental Figure S5.**
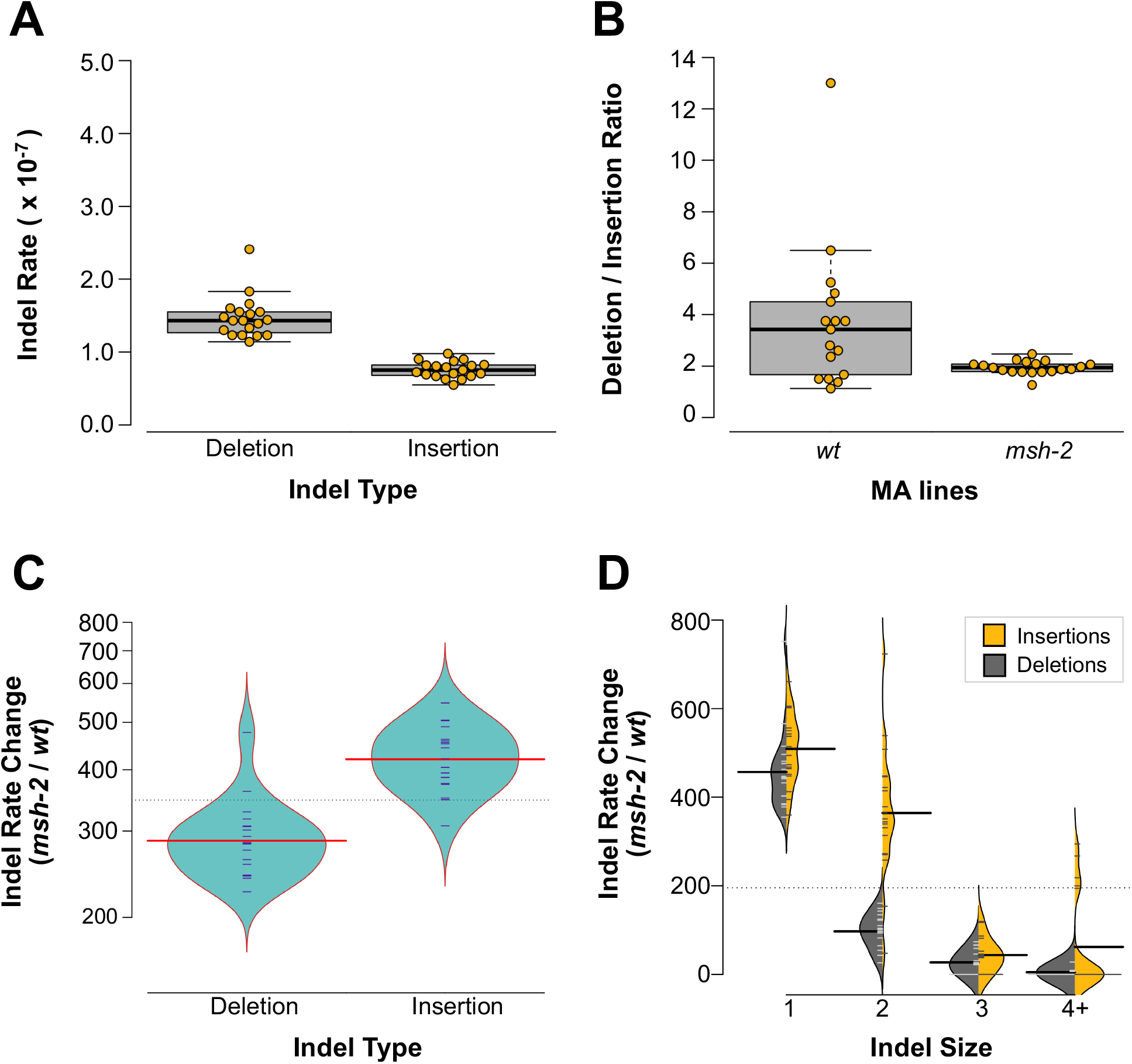
**Small indel rates in *msh-2* knockdown MA lines compared to wildtyp**e **MA lines** *(A)* The deletion rate in the *msh-2* knockdown lines was significantly greater than the insertion rate. *(B) msh-2* knockdown resulted in a significantly lower ratio of deletions per insertion relative to the wildtype MA lines. *(C)* The increase in the rate of insertions in *msh-2* knockdown relative to wildtype MA lines is significantly greater than the increase in the rate of deletions. *(D)* Both indel type (insertion *vs.* deletion), indel size, and the interaction of indel type and size have significant effects on the increase in indel rate in *msh-2* knockdown relative to wildtype MA lines. The rate of single base pair indels exhibits the greatest increase in *msh-2* knockdown MA lines, followed by 2-bp indels. Insertions increase more than deletions, especially the 2-bp insertions.

**Supplemental Figure S6.**
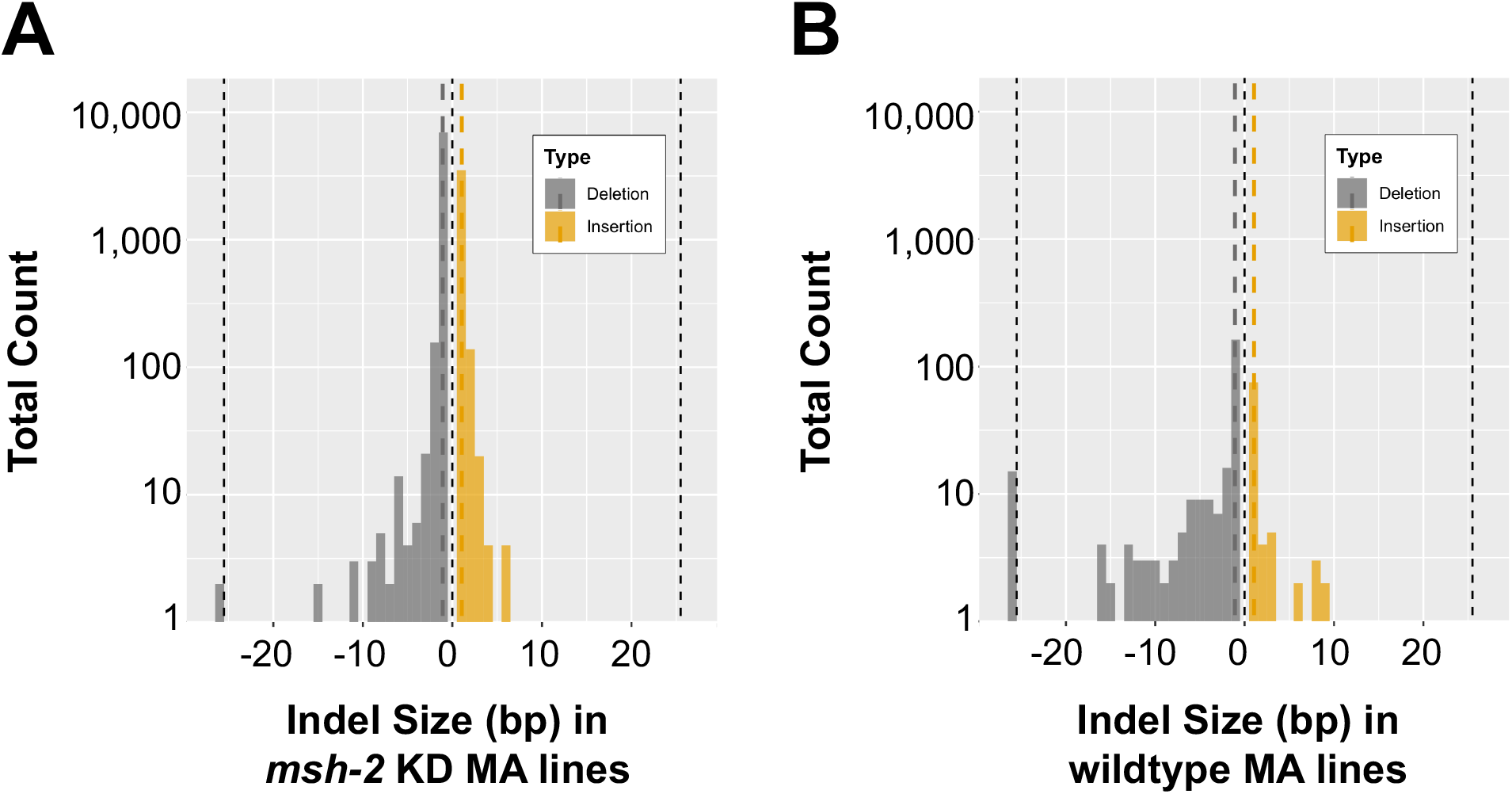
The size distribution of small indels. *(A)* The size distributions of both deletions and insertions in *msh-2* knockdown MA lines display a preponderance of single nucleotide indels. There was no difference in the average size of insertions (1.06 bp) and deletions (1.08 bp) in the *msh-2* knockdown lines. *(B)* In contrast, the average deletion size (6.38 bp) was significantly greater than the average insertion size (1.94 bp) in the wildtype MA lines.

**Supplemental Figure S7.**
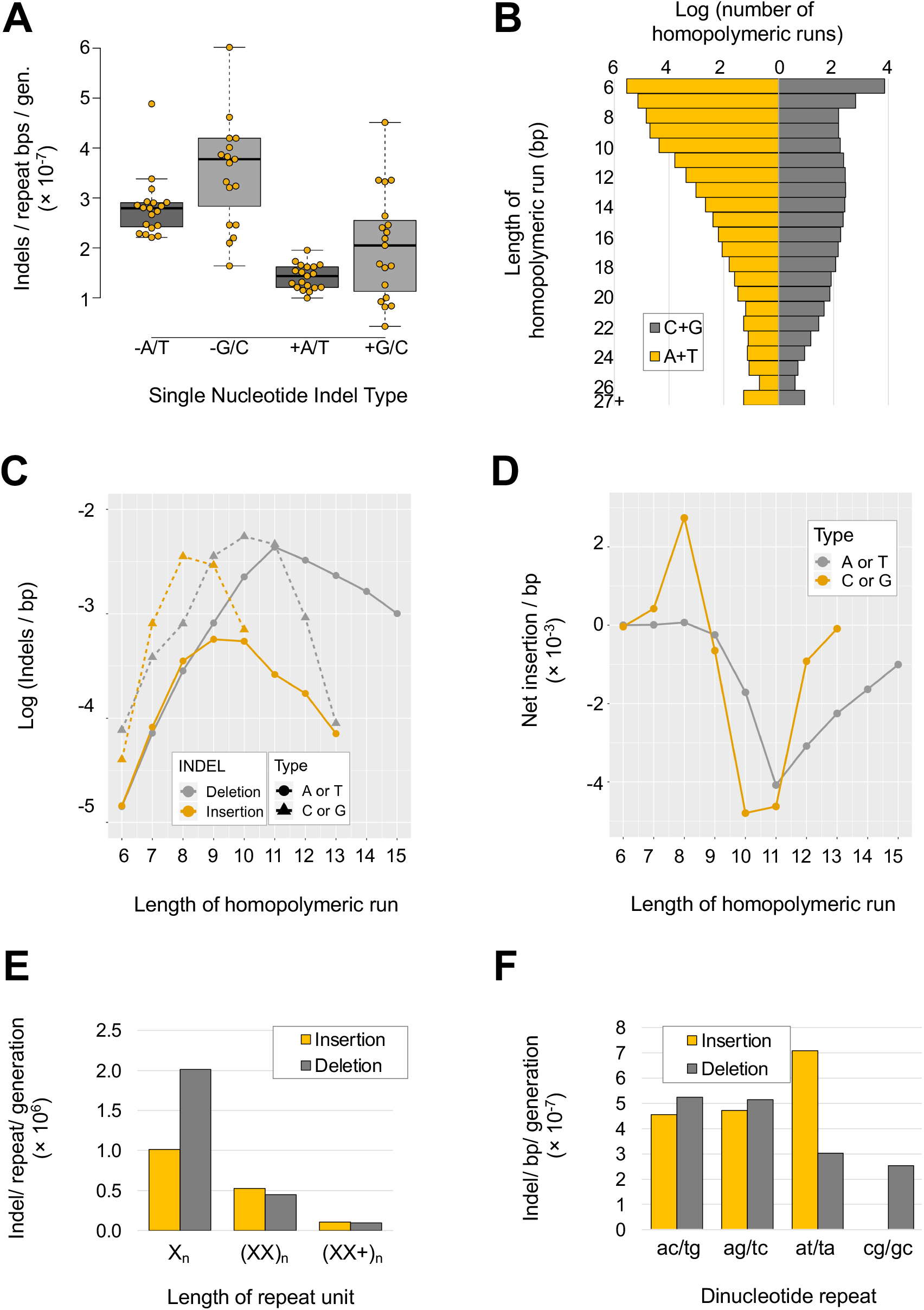
Variation in indel frequencies within indel repeats. *(A)* G/C homopolymeric runs of ≥6 bps in length have a higher indel rate per base per generation than A/T runs. Both indel type (deletions *vs.* insertions) and nucleotide (A+T versus C+G), have significant effects on the per base indel rate in homopolymeric runs. *(B)* The frequency and length distribution of homopolymeric runs depends on the type of nucleotide comprising the homopolymer. *(C)* The indel rates in homopolymeric runs vary with the length and type of nucleotide in the runs. Both insertion and deletion rates increase with the length of a run up to a point, and subsequently decline. The deletion rates were highest in runs of 10-11 bps for both A/T and G/C and lower for runs that were either smaller or larger. The insertion rates were highest when the length of a run was 8-9 bps for G/C and 8-10 bps for A/T. *(D)* The net gain and loss rates in homopolymeric runs vary with length and composition. The greatest net loss was in runs of 10-11 bps for both A/T and G/C. The greatest net gain was for G/C runs of 8 bps. *(E)* Deletions per repeat are only significantly higher than insertions in homopolymeric runs. In di- and polynucleotide repeats, insertions are roughly equal in frequency. *(F)* While most di-nucleotide repeats had a slightly higher deletion rate than insertion rate, AT dinucleotide repeats had an insertion rate that was almost twice as high as their deletion rate.

**Supplemental Figure S8.**
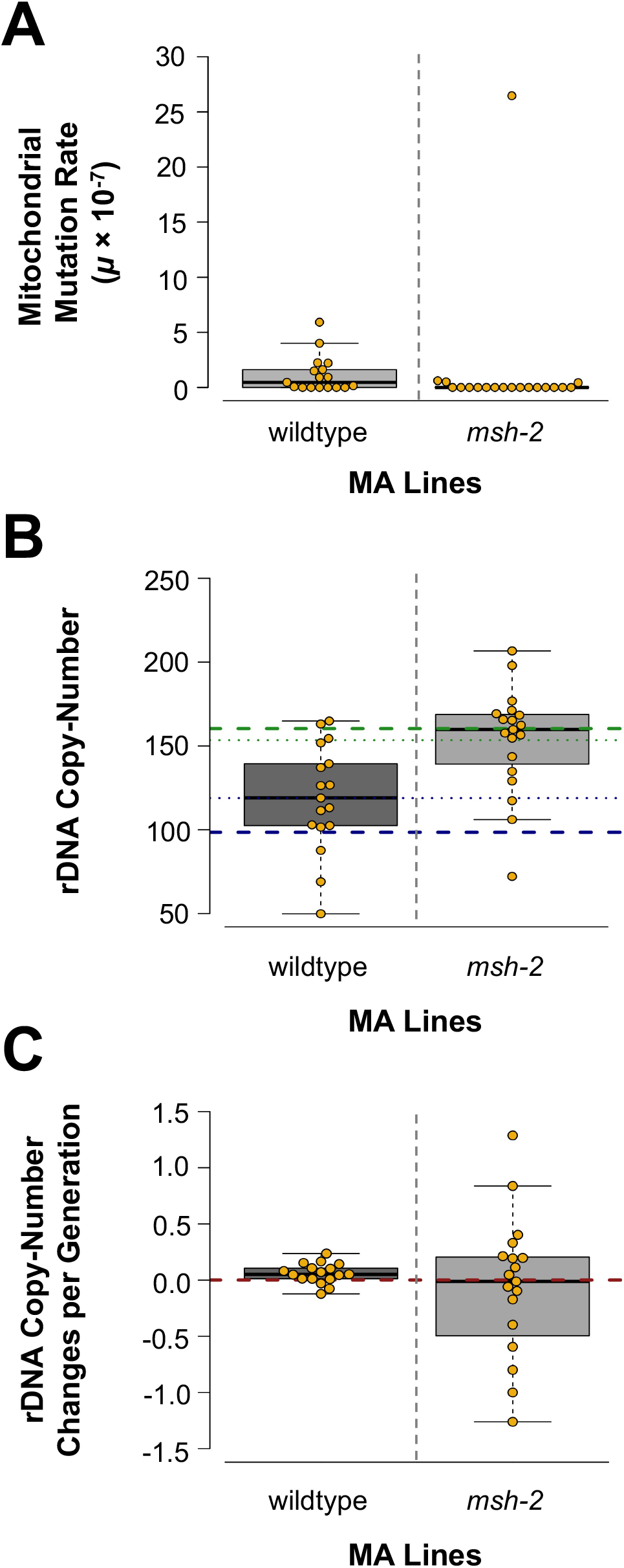
Comparisons between the *msh-2* knockdown and wildtype MA lines with respect to mitochondrial mutations, and changes rDNA copy-number. *(A)* Line-specific mtDNA mutation rate in the *msh-2* knockdown versus wildtype MA experiments. Due to the large number of *msh-2* knockdown MA lines with no mitochondrial mutations and a substantial difference in the number of generations, a statistical test of the difference in mtDNA mutation rates between wildtype and *msh-2* knockdown MA lines was not performed. rDNA copy-number increased by an average 20 copies in the wildtype MA lines whereas *msh-2* knockdown MA lines exhibited no significant change in rDNA copy-number relative to their ancestral state. The blue and green dashed line displays the estimated rDNA copy-number in the N2 wildtype ancestral strain and *msh-2* ancestral strain, respectively. *(C)* The variation in the rDNA copy-number change per generation relative to the ancestral state is significantly greater in the *msh-2* knockdown than in the wildtype MA lines.

**Supplemental Figure S9.**
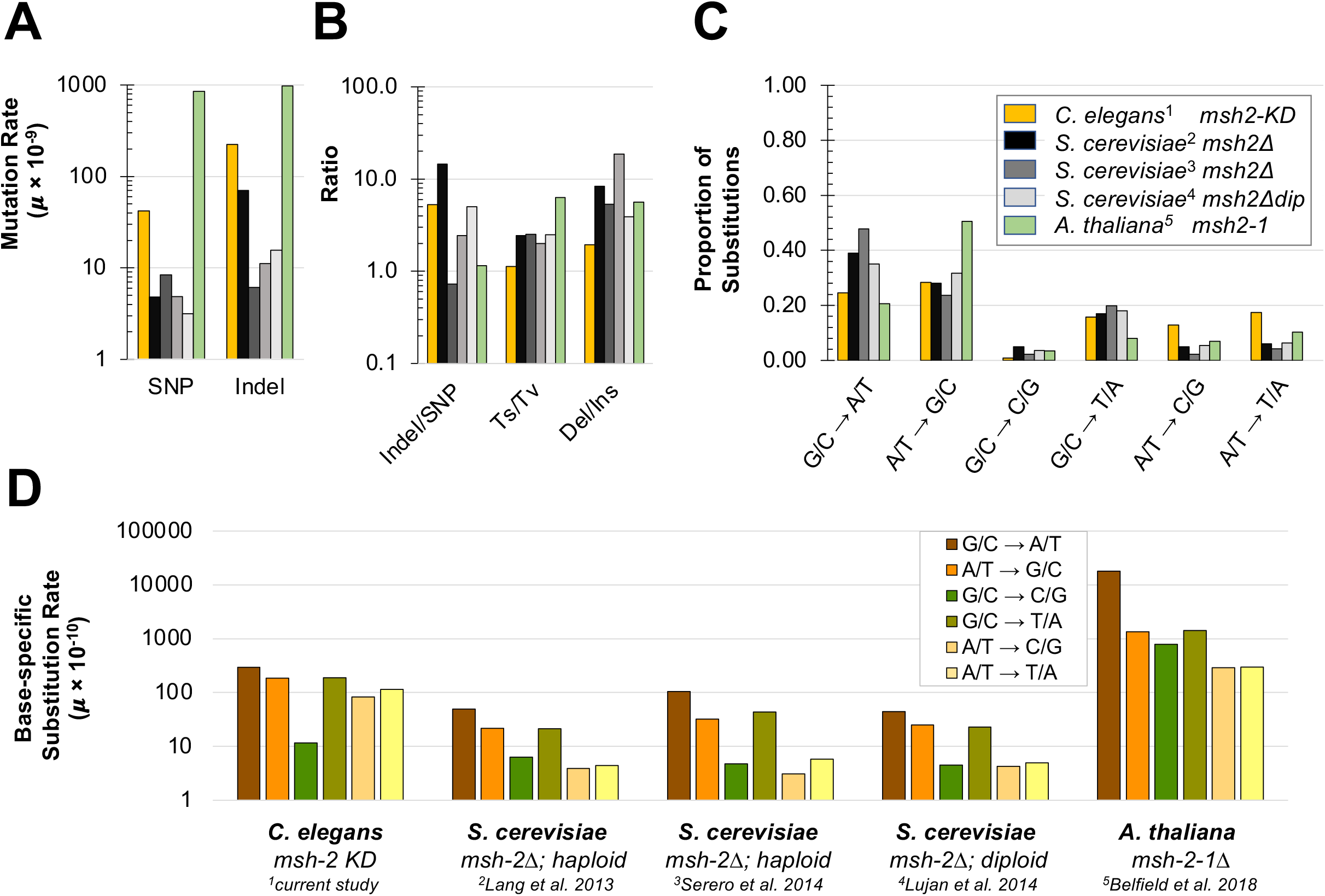
Comparisons of mutation rate and spectrum in mismatch repair-deficient lines of several model eukaryotic species. The mutational spectrum of the *Caenorhabditis elegans msh-2* knockdown lines analyzed in the present study^1^ are compared to knockout mutations of either *msh-2* or *msh-6* of the MutS*α* heterodimer in haploid (^2^Lang et al. 2013; ^3^Serero et al. 2014) and diploid *Saccharomyces cerevisiae* (^4^Lujan et al. 2014), and *Arabidopsis thaliana* (^5^Belfield et al. 2018). *(A)* SNP and indel rates estimated for the different datasets span three orders of magnitude. Substitution and indel rates in *A. thaliana* are approximately equal, while yeast species (except ^3^Serero et al. 2014) and *C. elegans* show markedly lower substitution rates than their respective indel rates. *(B)* The number of indels per SNP varies widely between taxa and between experiments with *Saccharomyces*. Transition/transversion (Ts/Tv) ratios are highest in *A. thaliana* and lowest for *C. elegans*. Deletions outnumber insertions in all of these species and the deletion/insertion ratio is lowest in *C. elegans*. *(C)* The mutational spectrum differs between species. *A thaliana* has a greater share of G/C → A/T transitions and lower share of A/T → G/C transitions relative to the other species. *C. elegans* has a greater share of A/T → C/G or T/A transversions relative to other species. *(D)* The base-specific substitution rates vary widely between different taxa.

## LITERATURE CITED

Alani E. 1996. The *Saccharomyces cerevisiae* Msh2 and Msh6 proteins form a complex that specifically binds to duplex oligonucleotides containing mismatched DNA base pairs. Mol Cell Biol. 16:5604–5615.

Antequera F, Tamame M, Villanueva J, Santos T. 1984. DNA methylation in the fungi. J Biol Chem. 259:8033–8036.

Aquilina G, Bignami M. 2001. Mismatch repair in correction of replication errors and processing of DNA damage. J Cell Physiol. 187:145–154.

Belfield EJ, Ding ZJ, Jamieson FJC, Visscher AM, Zheng SJ, Mithani A, Harberd NP. 2018. DNA mismatch repair preferentially protects genes from mutation. Genome Res. 28:66–74.

Bik HM, Fournier D, Sung W, Bergeron RD, Thomas WK. 2013. Intra-genomic variation in the ribosomal repeats of nematodes. PLoS One 8:e78230.

*C. elegans* Sequencing Consortium. 1998. Genome sequence of the nematode *C. elegans*: a platform for investigating biology. Science 282:2012–2018.

Culligan KM, Hays JB. 2000. *Arabidopsis* MutS homologs-AtMSH2, AtMSH3, AtMSH6, and a novel AtMSH7-form three distinct protein heterodimers with different specificities for mismatched DNA. Plant Cell 12:991–1002.

Danecek P, Auton A, Abecasis G, Albers CA, Banks E, DePristo MA, Handsaker RE, Lunter G, Marth GT, Sherry ST, et al. 2011. The variant call format and VCFtools. Bioinformatics 27:2156–2158.

Degtyareva NP, Greenwell P, Hofmann ER, Hengartner MO, Zhang L, Culotti JG, Petes TD. 2002. *Caenorhabditis elegans* DNA mismatch repair gene *msh-2* is required for microsatellite stability and maintenance of genome integrity. Proc Natl Acad Sci U S A. 99:2158–2163.

Denver DR, Feinberg S, Estes S, Thomas WK, Lynch M. 2005. Mutation rates, spectra and hotspots in mismatch repair-deficient *Caenorhabditis elegans*. Genetics 170:107–113.

Denver DR, Feinberg S, Steding C, Durbin M, Lynch M. 2006. The relative roles of three DNA repair pathways in preventing *Caenorhabditis elegans* mutation accumulation. Genetics 174:57–65.

Dillon MM, Sung W, Sebra R, Lynch M, Cooper VS. 2017. Genome-wide biases in the rate andmolecular spectrum of spontaneous mutations in *Vibrio cholerae* and *Vibrio fischeri*. Mol Biol Evol. 34:93–109.

Eisen JA, Hanawalt PC. 1999. A phylogenomic study of DNA repair genes, proteins, and processes. Mut Res. 435:171–213.

Evans KJ, Huang N, Stempor P, Chesney MA, Down TA, Ahringer J. 2016. Stable *Caenorhabditis elegans* chromatin domains separate broadly expressed and developmentally regulated genes. Proc Natl Acad Sci U S A. 113:E7020–E7029.

Falconer DS. 1989. Introduction to Quantitative Genetics. John Wiley & Sons, Inc., New York, NY.

Farslow JC, Lipinski KJ, Packard LB, Edgley ML, Taylor J, Flibotte S, Moerman DG, Katju V, Bergthorsson U. 2015. Rapid increase in frequency of gene copy-number variants during experimental evolution in *Caenorhabditis elegans*. BMC Genomics 16:1044.

Finnegan EJ, Peacock WJ, Dennis ES. 1996. Reduced DNA methylation in *Arabidopsis thaliana* results in abnormal plant development. Proc Natl Acad Sci U S A. 93:8449–8454.

Friedman J, Hastie, T, Tibshirani R. 2010. Regularization paths for generalized linear models via coordinate descent. J Stat Softw. 33:1–22.

Frøkjær-Jensen C, Jain N, Hansen L, Davis MW, Li Y, Zhao D, Rebora K, Millet JRM, Liu X, Kim SK, et al. 2016. An abundant class of non-coding DNA can prevent stochastic gene silencing in the *C. elegans* germline. Cell 166:343–357.

Garrison E, Marth G. 2012. Haplotype-based variant detection from short-read sequencing. arXiv preprint arXiv:1207.3907 [q-bio.GN].

Gibbons JG, Branco AT, Yu S, Lemos B. 2014. Ribosomal DNA copy number is coupled with gene expression variation and mitochondrial abundance in humans. Nat Commun. 5:4850.

Gómez R, Spampinato CP. 2013. Mismatch recognition function of *Arabidopsis thaliana* MutSγ. DNA Repair 12:257–264.

Gragg H, Harfe BD, Jinks-Robertson S. 2002. Base composition of mononucleotide runs affects DNA polymerase slippage and removal of frameshift intermediates by mismatch repair in *Saccharomyces cerevisiae*. Mol Cell Biol. 22:8756–8762.

Groothuizen FS, Sixma TK. 2016. The conserved molecular machinery in DNA mismatch repair enzyme structures. DNA Repair 38:14–23.

Habraken Y, Sung P, Prakash L, Prakash S. 1996. Binding of insertion/deletion DNA mismatches by the heterodimer of yeast mismatch repair proteins MSH2 and MSH3. Curr Biol. 6:1185–1187.

Harfe BD, Jinks-Robertson S. 2000. DNA mismatch repair and genetic instability. Annu Rev Genet. 34:359–399.

Harrington JM, Kolodner RD. 2007. *Saccharomyces cerevisiae* Msh2-Msh3 acts in repair of base-base mispairs. Mol Cell Biol. 27:6546–6554.

Harris TW, Antoshechkin I, Bieri T, Blasiar D, Chan J, Chen WJ, De La Cruz N, Davis P, Duesbury M, Fang R, et al. 2010. WormBase: a comprehensive resource for nematode research. Nucleic Acids Res. 38 (Database issue):D463–D467.

Jiricny J. 2006. The multifaceted mismatch-repair system. Nat Rev Mol Cell Biol. 7:335–346.

Jónsson H, Sulem P, Kehr B, Kristmundsdottir S, Zink F, Hjartarson E, Hardarson MT, Hjorleifsson KE, Eggertsson HP, Gudjonssonet SA, et al. 2017. Parental influence on human germline de novo mutations in 1,548 trios from Iceland. Nature 549:519–522.

Katju V, Bergthorsson U. 2019. Old trade, new tricks: insights into the spontaneous mutation process from the partnering of classical mutation accumulation experiments with high-throughput genomic approaches. Genome Biol Evol. 11:136–165.

Katju V, LaBeau EM, Lipinski KJ, Bergthorsson U. 2008. Sex change by gene conversion in a *Caenorhabditis elegans fog-2* mutant. Genetics 180:669–672.

Katju V, Packard LB, Bu L, Keightley PD, Bergthorsson U. 2015. Fitness decline in spontaneous mutation accumulation lines of *Caenorhabditis elegans* with varying effective population sizes. Evolution 69:104–116.

Katju V, Packard LB, Keightley PD. 2018. Fitness decline under osmotic stress in *Caenorhabditis elegans* populations subjected to spontaneous mutation accumulation at varying population sizes. Evolution 72:1000–1008.

Kamath RS, Martinez-Campos M, Zipperlein P, Frazer AG, Ahringer J. 2001 Effectiveness of specific RNA-mediated interference through ingested double-stranded RNA in *Caenorhabditis elegans*. Genome Biol. 2:research0002.

Keightley PD, Trivedi U, Thomson M, Oliver F, Kumar S, Blaxter ML. 2009. Analysis of the genome sequences of three *Drosophila melanogaster* spontaneous mutation accumulation lines. Genome Res. 19:1195–1201.

Kolodner R. 1996. Biochemistry and genetics of eukaryotic mismatch repair. Genes Dev 10:1433–1442.

Konrad A, Brady MJ, Bergthorsson U, Katju V. 2019. Mutational landscape of spontaneous base substitutions and small indels in experimental *Caenorhabditis elegans* populations of differing size. Genetics 212:837–854.

Konrad A, Flibotte S, Taylor J, Waterston RH, Moerman DG, Bergthorsson U, Katju V. 2018. Mutational and transcriptional landscape of spontaneous gene duplications and deletions in *Caenorhabditis elegans*. Proc Natl Acad Sci U S A. 115:7386–7391.

Konrad A, Thompson O, Waterston RH, Moerman DG, Keightley PD, Bergthorsson U, Katju V. 2017. Mitochondrial mutation rate, spectrum and heteroplasmy in *Caenorhabditis elegans* spontaneous mutation accumulation lines of differing size. Mol Biol Evol. l34:1319–1334.

Kunkel TA, Erie DA. 2005. DNA mismatch repair. Annu Rev Biochem. 74:681–710.

Lang GI, Parsons L, Gammie AE. 2013. Mutation rates, spectra, and genome-wide distribution of spontaneous mutations in mismatch repair deficient yeast. G3 3:1453–1465.

Lee H, Popodi E, Tanga H, Foster PL. 2012. Rate and molecular spectrum of spontaneous mutations in the bacterium *Escherichia coli* as determined by whole-genome sequencing. Proc Natl Acad Sci U S A. 109:E2774–2783.

Lee SD, Surtees JA, Alani E. 2007. *Saccharomyces cerevisiae* MSH2-MSH3 and MSH2-MSH6 complexes display distinct requirements for DNA binding domain I in mismatch recognition. J Mol Biol. 366:53–66.

Li H. 2011. Improving SNP discovery by base alignment quality. Bioinformatics 27:1157–1158.

Li H., Durbin R. 2009. Fast and accurate short read alignment with Burrows-Wheeler transform. Bioinformatics 25:1754–1760.

Li H, Handsaker B, Wysoker A, Fennell T, Ruan J, Homer N, Marth G, Abecasis G, Durbin R, 1000 Genome Project Data Processing Subgroup. 2009. The Sequence alignment/map (SAM) format and SAMtools. Bioinformatics 25:2078–2079.

Lin Z, Nei M, Ma H. 2007. The origins and early evolution of DNA mismatch repair genes— multiple horizontal gene transfers and co-evolution. Nucleic Acids Res. 35:7591–7603.

Lindahl T. 1993. Instability and decay of the primary structure of DNA. Nature 362:709–715.

Long H, Miller SF, Strauss C, Zhao C, Cheng L, Ye Z, Griffin K, Te R, Lee H, Chen C-C, et al. 2016. Antibiotic treatment enhances the genome-wide mutation rate of target cells. Proc Natl Acad Sci U S A. 113:E2498–E2505.

Long H, S. Miller SF, Williams E, Lynch M. 2018. Specificity of the DNA mismatch repair system (MMR) and mutagenesis bias in bacteria. Mol Biol Evol. 35:2414–2421.

Lutsenko E, Bhagwat AS. 1999. Principal causes of hot spots for cytosine to thymine mutations at sites of cytosine methylation in growing cells. A model, its experimental support and implications. Mutat Res. 437:11–20.

Lynch HT, Lynch PM, Lanspa SJ, Snyder CL, Lynch JF, Boland CR. 2009. Review of the Lynch syndrome: history, molecular genetics, screening, differential diagnosis, and medicolegal ramifications. Clin Genet. 76:1–18.

Maydan JS, Flibotte S, Edgley ML, Lau J, Selzer RR, Richmond TA, Pofahl NJ, Thomas JH, Moerman DG. 2007. Efficient high-resolution deletion discovery in *Caenorhabditis elegans* by array comparative genomic hybridization. Genome Res. 17:337–347.

Maydan JS, Lorch A, Edgley ML, Flibotte S, Moerman DG. 2010. Copy number variation in the genomes of twelve natural isolates of *Caenorhabditis elegans*. BMC Genomics 11:62.

Meier B, Volkova NV, Hong Y, Schofield P, Campbell PJ, Gerstung M, Gartner A. 2018. Mutational signatures of DNA mismatch repair deficiency in *C. elegans* and human cancers. Genome Res. 28:666–675.

Miyata T, Hayashida H, Kuma K, Mitsuyasu K, Yasunaga T. 1987. Male-driven molecular evolution: a model and nucleotide sequence analysis. Cold Spring Harbor Symp Quant Biol. 52:863–867

Modrich P. 1991. Mechanisms and biological effects of mismatch repair. Annu Rev Genet. 25:229–253.

Morgulis A, Gertz EM, Schäffer AA, Agarwala R. 2006. A fast and symmetric DUST implementation to mask low-complexity DNA sequences. J Comput Biol. 13:1028–1040.

Mudunuri SB, Nagarajaram HA. 2007. IMEx: Imperfect Microsatellite Extractor. Bioinformatics 23:1181–1187.

Ness RW, Morgan AD, Vasanthakrishnan RB, Colegrave N, Keightley PD. 2015. Extensive de novo mutation rate variation between individuals and across the genome of *Chlamydomonas reinhardtii*. Genome Res. 25:1739–1749.

Paredes S, Maggert KA. 2009. Ribosomal DNA contributes to global chromatin regulation. Proc Natl Acad Sci U S A. 106:17829–17834.

R Core Development Team. 2014. R: A language and environment for statistical computing. R Foundation for Statistical Computing, Vienna, Austria. URL http://www.R-project.org/.

Ratan A, Olson TL, Loughran TP, Miller W. 2015. Identification of indels in next-generation sequencing data. BMC Bioinformatics 16:42.

Richman S. 2015. Deficient mismatch repair: Read all about it. Int J Oncol. 47:1189–1202.

Rimmer A, Phan H, Mathieson I, Iqbal Z, Twigg SRF, WGS500 Consortium; Wilkie AOM, McVean G, Lunter G. 2014. Integrating mapping-, assembly- and haplotype-based approaches for calling variants in clinical sequencing applications. Nat Genet. 46:912–918.

Rockman MV, Kruglyak L. 2009. Recombinational landscape and population genomics of *Caenorhabditis elegans*. PLoS Genet. 5:e1000419.

Sachadyn P. 2010. Conservation and diversity of MutS proteins. Mut Res. 694:20–30.

Saxena AS, Salomon MP, Matsuba C, Yeh S-D, Baer CF. 2019. Evolution of the mutational process under relaxed selection in *Caenorhabditis elegans*. Mol Biol Evol. 36:239–251.

Schedl T, Kimble J. 1988. *fog-2*, a germ-line-specific sex determination gene required for hermaphrodite spermatogenesis in *Caenorhabditis elegans*. Genetics 119:43–61.

Serero A, Jubin C, Loeillet S, Legoix-Né P, Nicolas AG. 2014. Mutational landscape of yeast mutator strains. Proc Natl Acad Sci U S A. 111:1897–1902.

Shodhan A, Lukaszewicz A, Novatchkova M, Loidl J. 2014. Msh4 and Msh5 function in SC-independent chiasma formation during the streamlined meiosis of *Tetrahymena*. Genetics 198:983–993.

Simpson VJ, Johnson TE, Hammen RF. 1986. *Caenorhabditis elegans* DNA does not contain 5-methylcytosine at any time during development or aging. Nucleic Acids Res. 14:6711–6719.

Strand M, Prolla TA, Liskay RM, Petes TD. 1993. Destabilization of tracts of simple repetitive DNA in yeast by mutations affecting DNA mismatch repair. Nature 365:274–276.

Streisinger G, OkadaY, Emrich J, Newton J, Tsugita A, Terzaghi E, Inouye M. 1966. Frameshift mutations and the genetic code. Cold Spring Harbor Symp Quant Biol. 31:77–84.

Sun L, Zhang Y, Zhang Z, Zheng Y, Du L, Zhu B. 2016. Preferential protection of genetic fidelity within open chromatin by the mismatch repair machinery. J Biol Chem. 291:17692– 17705.

Sung W, Ackerman MS, Gout JF, Miller SF, Williams E, Foster PL, Lynch M. 2015. Asymmetric context-dependent mutation patterns revealed through mutation-accumulation experiments. Mol Biol Evol. 32:1672–1683.

Takemoto N, Numata I, Su’etsugu M, Miyoshi-Akiyama T. 2018. Bacterial EndoMS/NucS acts as a clamp-mediated mismatch endonuclease to prevent asymmetric accumulation of replication errors. Nucleic Acids Res. 46:6152–6165.

Takuno S, Ran J-H, Gaut BS. 2016. Evolutionary patterns of genetic DNA methylation vary across land plants. Nat Plants. 2:15222.

Tang Y, Gao XD, Wang Y, Yuan BF, Feng YQ. 2012. Widespread existence of cytosine methylation in yeast DNA measured by gas chromatography/mass spectrometry. Anal Chem. 84:7249–7255.

Thompson O, Edgley M, Strasbourger P, Flibotte S, Ewing B, Adair R, Au V, Chaudhry I, Fernando L, Hutter H, et al. 2013. The million mutation project: a new approach to genetics in *Caenorhabditis elegans*. Genome Res. 23:1749–1762.

Tijsterman M, Pothof J, Plasterk RH. 2002. Frequent germline mutations and somatic repeat instability in DNA mismatch-repair-deficient *Caenorhabditis elegans*. Genetics 161:651– 660.

Wang X, Zhao Y, Wong K, Ehlers P, Kohara Y, Jones SJ, Marra MA, Holt RA, Moerman DG, Hansen D. 2009. Identification of genes expressed in the hermaphrodite germ line of *C. elegans* using SAGE. BMC Genomics 10:213.

Whittle C-A, Johnston MO. 2003. Male-biased transmission of deleterious mutations to the progeny in *Arabidopsis thaliana*. Proc Natl Acad Sci U S A. 100:4055–4059.

Wilson Sayres MA, Makova KD. 2011. Genome analyses substantiate male mutation bias in many species. Bioessays 33:938–945.

Wright S. 1931. Evolution in Mendelian populations. Genetics 16:97–159.

Yang Y, Karthikeyan R, Mack SE, Vonarx EJ, Kunz BA. 1999. Analysis of yeast *pms1*, *msh2*, and *mlh1* mutators points to differences in mismatch correction efficiencies between prokaryotic and eukaryotic cells. Mol Gen Genet. 261:777–787.

Zalevsky J, MacQueen AJ, Duffy JB, Kemphues KJ, Villeneuve AM. 1999. Crossing over during *Caenorhabditis elegans* meiosis requires a conserved MutS-based pathway that is partially dispensable in budding yeast. Genetics 153:1271–1283.

